# Structure of VanS from Vancomycin-Resistant Enterococci: A Sensor Kinase with Weak ATP Binding

**DOI:** 10.1101/2022.12.12.520123

**Authors:** Kimberly C. Grasty, Claudia Guzik, Elizabeth J. D’Lauro, Shae B. Padrick, Joris Beld, Patrick J. Loll

## Abstract

The VanRS two-component system regulates the resistance phenotype of vancomycin-resistant enterococci (VRE). VanS is a sensor histidine kinase that responds to the presence of vancomycin by autophosphorylating and subsequently transferring the phosphoryl group to the response regulator, VanR. The phosphotransfer activates VanR as a transcription factor, which initiates the expression of resistance genes. Structural information about VanS proteins has remained elusive, hindering the molecular-level understanding of their function. Here, we present X-ray crystal structures for the catalytic and ATP-binding (CA) domains of two VanS proteins, derived from VRE types A and C. Both proteins adopt the canonical Bergerat fold that has been observed for CA domains of other prokaryotic histidine kinases. We attempted to determine structures for the nucleotide-bound forms of both proteins; however, despite repeated efforts, these forms could not be crystallized, prompting us to measure the proteins’ binding affinities for ATP. Unexpectedly, both CA domains displayed low affinities for the nucleotide, with *K*_D_ values in the low millimolar range. Since these *K*_D_ values are comparable to intracellular ATP concentrations, this weak substrate binding could reflect a way of regulating expression of the resistance phenotype.

## Introduction

For many years after its introduction, vancomycin occupied an important niche as a last-resort antibiotic, reserved for patients with beta-lactam allergies or infections refractory to other antibiotics. However, in the latter part of the 20^th^ century, antibiotic-resistant pathogens such as methicillin-resistant *Staphylococcus aureus* (MRSA) began spreading rapidly, which elicited wider usage of vancomycin (1). Inevitably, this increased use led to increased levels of resistance. In particular, vancomycin resistance has now become commonplace in enterococci, transforming these organisms from largely benign commensals to <u>v</u>ancomycin-<u>r</u>esistant <u>e</u>nterococci (VRE), serious pathogens that have been flagged by the World Health Organization as high-priority targets for therapeutic development (2,3). VRE are now counted among the so-called ESKAPE pathogens, which are responsible for the majority of hospital-acquired infections and can be very challenging to treat (4).

Vancomycin interferes with production of the bacterial cell wall. It does so by binding the D-Ala-D-Ala sequence found at the C-terminus of Lipid II’s muramyl peptide, thereby sequestering a critical precursor in peptidoglycan biosynthesis (5,6). Bacteria become resistant to vancomycin by acquiring a suite of genes that encode various remodeling enzymes, which degrade D-Ala-D-Ala and replace it with D-Ala-D-Ser or D-Ala-D-Lac, neither of which is efficiently recognized by the antibiotic (7). Expression of the remodeling-enzyme genes is controlled by VanS and VanR, which form a two-component system that senses the presence of the antibiotic and transduces this signal (8). Multiple VRE genotypes have been identified, corresponding to the acquisition of similar but distinct vancomycin-resistance genes; these genotypes are denoted as VRE types A through N (9).

Two-component systems are ubiquitous in bacteria, fungi, and plants (10-12), and consist of a sensor and a response regulator. The sensor (VanS in this case) is typically a transmembrane histidine kinase that autophosphorylates on a conserved histidine residue in response to an appropriate signal; the phosphoryl group is then transferred to the response regulator (here, VanR). VanR is a transcription factor, and phosphorylation enhances its affinity for the promoter controlling expression of the remodeling genes (13). Thus, in the presence of vancomycin, VanS will autophosphorylate and subsequently transfer the phosphoryl group to VanR, thereby activating transcription of the genes encoding the remodeling enzymes.

The VanS sensor kinase is organized into several structural domains (Figure 1A), which include a periplasmic sensor domain flanked by two transmembrane helices, a membrane-proximal HAMP domain, a <u>d</u>imerization-and-<u>h</u>istidine-<u>p</u>hosphorylation (DHp) domain, and a <u>c</u>atalytic <u>A</u>TP-binding (CA) domain (14-16). The latter plays a dominant role in the autophosphorylation mechanism, as it contributes all of the residues responsible for proper positioning of the nucleotide during catalysis (17). Since CA domains are essential for histidine kinase activity (and therefore for two-component signaling), it is not surprising that these domains are being actively explored as potential targets for antimicrobial therapeutics (18).

**Figure 1.**
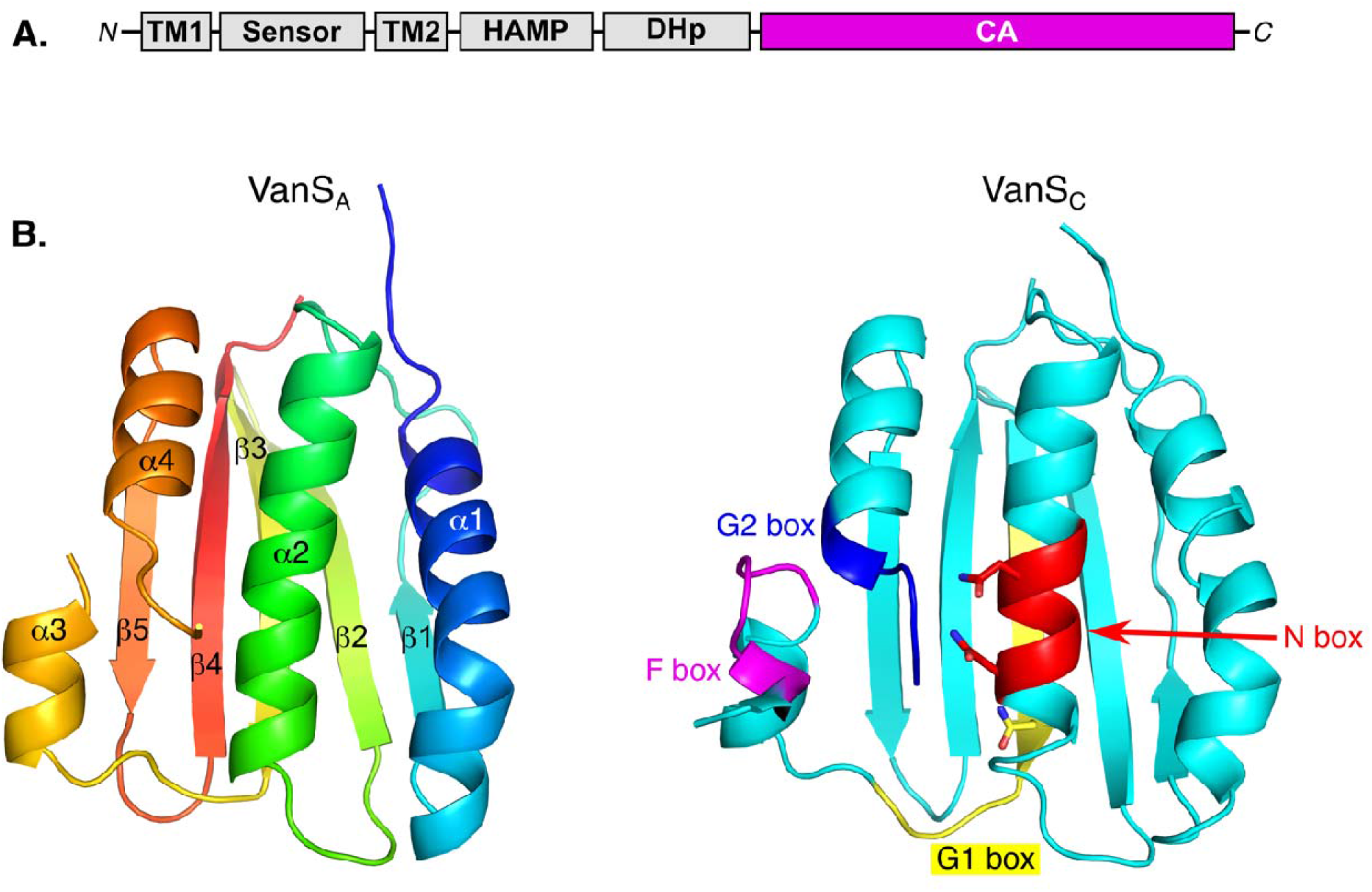
Structures of the VanS_A_ and VanS_C_ CA domains. (A) Schematic illustration of the VanS domain structure. (B) Ribbon diagram showing the VanS_A_ CA-domain structure, colored using “rainbow” scheme that ranges from blue at the N-terminus to red at the C-terminus. The numbering of th secondary-structure elements is also illustrated on the VanS_A_ structure (this numbering is the same for both VanS_A_ and VanS_C_). (C) The VanS_C_ CA-domain structure, with key structural motifs highlighted (these motifs are present in both VanS_A_ and VanS_C_). These motifs include the N box (red; the two conserved Asn residues are shown), the G1 box (yellow, including the non-canonical Asn residue found in both VanS_A_ and VanS_C_), the F box (magenta), and the G2 box (blue). Stereo versions of panels (B) & (C) can be found in Supporting Information Figure S9.

In the VanRS two-component system, structures are available for the inactive and activated forms of VanR (19), but comparable information is lacking for VanS. Thus, to gain further insight into the functioning of the VanRS system, we characterized the structures and functions of CA domains derived from two different VanS proteins. We report here the X-ray crystal structures of the VanS CA domains associated with type-A and type-C VRE. Unexpectedly, both CA domains were found to display unusually low affinities for their ATP substrate, raising intriguing questions about the potential contributions of nucleotide binding to the overall regulation of VanS activity.

## Results

### Structure determination of the VanS_A_ and VanS_C_ CA domains

As part of an effort to understand the molecular basis of vancomycin resistance in VRE, we purified and crystallized the CA domains of the VanS proteins associated with A- and C-type resistance, and determined their structures at resolutions of 2.2 and 1.45 Å respectively (Table 1; Supporting Information Figures S1-S3). We will refer to the type-A and type-C proteins as VanS_A_ and VanS_C_.

**Table 1.** Data collection and refinement statistics.

| Data Collection Statistics | VanS<sub>A</sub> 7 keV | VanS<sub>A</sub> 13.5 keV | VanS<sub>C</sub> |
| --- | --- | --- | --- |
| PDB ID | 8DWZ | 8DVQ | 8DX0 |
| Diffraction source | Beamline 24-ID-C, APS | Beamline 17-ID-1, NSLS-II | Beamline 24-ID-C, APS |
| Wavelength (Å) | 1.7712 | 0.92009 | 0.9791 |
| Temperature (K) | 100 | 100 | 100 |
| Detector | Dectris Pilatus 6MF | Eiger 9M | Dectris Pilatus 6MF |
| Resolution range (Å)<sup>a</sup> | 56.1 – 2.21<br>(2.29 – 2.21) | 28.4 – 2.19<br>(2.27 – 2.19) | 64.7 – 1.45<br>(1.50 – 1.45) |
| Spacegroup | *P*2<sub>1</sub>2<sub>1</sub>2 | *P*2<sub>1</sub>2<sub>1</sub>2 | *P*2<sub>1</sub> |
| Unit cell |  |  |  |
| *a*, *b*, *c* (Å) | 69.83, 94.40, 29.77 | 69.83, 94.40, 29.77 | 62.70, 40.23, 64.71 |
| $\alpha$, $\beta$, $\gamma$ (°) | 90.0, 90.0 90.0 | 90.0, 90.0 90.0 | 90.0, 91.64, 90.0 |
| Total number of observations | 176,601 (5296) | 91,282 (8083) | 345,447 (15,699) |
| Number of unique reflections | 9612 (664) | 10,676 (1031) | 55,912 (4842) |
| Average multiplicity | 18.4 (8.0) | 8.55 (7.84) | 6.2 (3.1) |
| Completeness (%) | 92.8 (63.6) | 99.7 (98.0) | 97.1 (84.7) |
| Mean I/sigma(I) | 23.1 (3.1) | 13.0 (2.2) | 22.8 (1.5) |
| Estimated Wilson B-factor (Å<sup>2</sup>) | 40.4 | 43.4 | 21.9 |
| R-merge<sup>b</sup> | 0.101 (0.578) | 0.090 (0.960) | 0.036 (0.600) |
| R-meas<sup>c</sup> | 0.104 (0.616) | 0.096 (1.027) | 0.039 (0.718) |
| R-pim<sup>d</sup> | 0.023 (0.206) | 0.033 (0.360) | 0.016 (0.386) |
| CC<sub>1/2</sub><sup>e</sup> | 0.999 (0.932) | 0.999 (0.849) | 0.999 (0.682) |
| CC<sub>1/2,anom</sub><sup>e</sup> | 0.65 (0.10) | --- | --- |
| Resolution at which CC<sub>1/2,anom</sub> drops below 0.3 (Å) | 2.6 | --- | --- |
| **Refinement and Model Statistics** |  |  |  |
| Number of reflections used | 17,544 (1807) | 10,620 (1029) | 51,113 (4534) |
| Reflections used for R-free | 911 (98) | 533 (45) | 2630 (223) |
| Rwork | 0.187 (0.268)<sup>f</sup> | 0.192 (0.273) | 0.185 (0.276) |
| Rfree | 0.234 (0.328)<sup>f</sup> | 0.239 (0.292) | 0.200 (0.312) |
| Number of non-hydrogen atoms |  |  |  |
| Protein | 1006 | 1015 | 2245 |
| Solvent | 24 | 45 | 228 |
| Ions | 2 | 2 | 3 |
| Average B-factor (Å) | 55.6 | 63.0 | 28.1 |
| RMS deviations from ideality |  |  |  |
| Bonds (Å) | 0.007 | 0.01 | 0.01 |
| Angles (°) | 0.84 | 1.27 | 1.07 |
| Residue distribution in Ramachandran plot |  |  |  |
| Most favored region (%) | 99.2 | 98.4 | 98.9 |
| Allowed (%) | 0.8 | 1.6 | 1.1 |
| Outliers (%) | 0 | 0 | 0 |
| Clashscore | 2.0 | 3.96 | 2.37 |

<sup>a</sup> Values in parentheses refer to the highest resolution shell.<sup>b</sup> $R_{\text{merge}}$ is calculated by the equation $R_{\text{merge}} = \sum_{hkl} \sum_i |I_i(hkl) - \langle I(hkl) \rangle| / \sum_{hkl} \sum_i I_i(hkl)$ , where $I_i(hkl)$ is the $i^{\text{th}}$ measurement.<sup>c</sup> $R_{\text{meas}}$ (or redundancy-independent $R_{\text{merge}}$ ) is calculated by the equation $R_{\text{meas}} = \sum_{hkl} [N(N-1)]^{1/2} \sum_i |I_i(hkl) - \langle I(hkl) \rangle| / \sum_{hkl} \sum_i I_i(hkl)$ , where $I_i(hkl)$ is the $i^{\text{th}}$ measurement and $N$ is the redundancy of each unique reflection $hkl$ (53).<sup>d</sup> $R_{\text{pim}}$ is calculated by the equation $R_{\text{pim}} = \sum_{hkl} [1/(N-1)]^{1/2} \sum_i |I_i(hkl) - \langle I(hkl) \rangle| / \sum_{hkl} \sum_i I_i(hkl)$ , where $I_i(hkl)$ is the $i^{\text{th}}$ measurement and $N$ is the redundancy of each unique reflection $hkl$ (54).<sup>e</sup> $CC_{1/2}$ is the correlation coefficient between two randomly chosen half data sets (55).<sup>f</sup> For the VanS<sub>A</sub> 7-keV data set, $F(+)$ and $F(-)$ were treated as separate observations during refinement.

The structure of the VanS_A_ CA domain was determined by the SAD method, relying upon two ordered cadmium ions contributed by the crystallization buffer. The crystal asymmetric unit contains one copy of the CA domain, and the cadmium ions mediate two of the three lattice contacts that connect different VanS_A_ molecules within the crystal (Figure S4). The structure was refined separately against two different data sets, one which was measured at low energy (7 keV) and used for phasing, while the other was collected at an energy more typically used for protein crystallography (13.5 keV). The structures obtained from the two different data sets are essentially identical, with a few minor differences in side-chain conformers, and have a root-mean square difference (RMSD) in Cα positions of 0.14 Å (0.59 Å for all non-hydrogen atoms).

The structure of the VanS_C_ CA domain was determined by molecular replacement, using an ensemble of homology models as probes. The crystal asymmetric unit contains two copies of the CA domain, related by an approximate two-fold axis of symmetry. The two protomers in the asymmetric unit are highly similar, with superposition yielding an RMSD for Cα positions of 0.25 Å (0.74 Å for all atoms). Metal ions also figure in the crystal packing of the VanS_C_ protein, with three Mg^2+^ ions being situated in two different lattice interfaces (Figure S5).

### Overall structure of the VanS CA domains

Both the VanS_A_ and VanS_C_ CA domains adopt the canonical Bergerat fold, which is associated with the GHKL superfamily of nucleotide-binding proteins, and in particular is characteristic of prokaryotic histidine kinases (14,15). Like other CA domains, the VanS_A_ and VanS_C_ structures feature four alpha helices packed against one face of a five-stranded antiparallel beta sheet (Figure 1). Also like other CA domains, the VanS structures contain a rare left-handed beta-alpha-beta crossover, corresponding to the β1-α2-β2 secondary structural elements. The presumptive nucleotide-binding site lies within a pocket bounded on either side by helices α2 and α4 and at the bottom by strands β3-5.

A series of conserved sequence motifs are associated with CA domains, namely the N, G1, F, and G2 boxes (20), and these motifs map to various positions clustered around the nucleotide-binding site. The N box is found on helix α2, and derives its name from the second asparagine in the NxxxNA consensus sequence. This asparagine is highly conserved and typically coordinates the metal ion of the bound ATP-Mg^2+^ complex; it corresponds to Asn-281 in VanS_A_ and Asn-264 in VanS_C_.

The G1 box is found at the C-terminus of strand β3, which runs directly under helix α2. The canonical G1 consensus sequence is DxGxG. The aspartate residue of this sequence typically forms a salt bridge with the amine group of the nucleotide’s adenine base, while the first conserved glycine marks the point where the polypeptide chain makes a sharp ∼90° turn to run underneath the nucleotide-binding pocket. Interestingly, neither VanS_A_ nor VanS_C_ conform to this consensus sequence, having asparagine residues in place of the conserved aspartate, and lacking the second glycine altogether. Both deviations from the consensus are rare, but not unprecedented. In particular, the Asp-to-Asn substitution is isosteric and, while it would eliminate the salt bridge, it should still allow for hydrogen bonding with the adenine base, and indeed the structure of the *T. maritima* kinase ThkA shows just such an asparagine-adenine hydrogen bond (21). The role of the second glycine in the G1 box is less clear, as it does not appear to contact the nucleotide in structures of CA domain-ATP complexes.

Immediately downstream of the G1 box lies an extended sequence that includes the short helix α3, followed by the other two conserved sequence motifs, the F box and the G2 box. This portion of the CA domain is known as the ATP lid, and is thought to close over the binding site once nucleotide is acquired. In the structures of both VanS_A_ and VanS_C_, a significant stretch of this extended region is disordered. It is common for part or all of the ATP lid to be disordered in structures of nucleotide-free CA domains; in some cases, partial disorder remains even when nucleotides are bound (e.g. PDB IDs 5C93 and 3SL2, corresponding to the kinases WalK and YycG). For both VanS_A_ and VanS_C_, the F-box is found just before the beginning of the disordered sequence, while the G2-box is found at the C-terminal end of the disordered region.

In addition to the motifs surrounding the nucleotide-binding site, CA domains also typically contain a conserved stretch of residues on the outward-facing surface of helix α4, immediately downstream of the G2-box. During autophosphorylation, these residues interact with the histidine kinase so as to properly position the CA domain, leading to α4 being known as the “Gripper” helix (22). The conserved residues are arrayed along the outer face of the helix and are generally hydrophobic. In VanS_A_ and VanS_C_, these so-called “sticky-finger” residues are (Ile-345, Ile-349, Gln-352) and (Ile-328, Ile-332, Ala-335), respectively.

### Comparison of the VanS_A_ and VanS_C_ CA domains

The sequences of the VanS_A_ and VanS_C_ CA are 37% identical, and both proteins adopt the same overall fold. To compare the two structures in more detail, we used TM-ALIGN to calculate a pairwise structural alignment (23). The VanS_A_ and VanS_C_ constructs contain 164 and 155 residues, respectively, and TM-ALIGN generated a superposition based upon 126 residues, with an RMSD value of 1.68 Å (Figure 2). Pairwise superposition using the protein LSQKAB yields an RMSD value of 0.14 Å for alpha-carbon positions alone (24).

**Figure 2.**
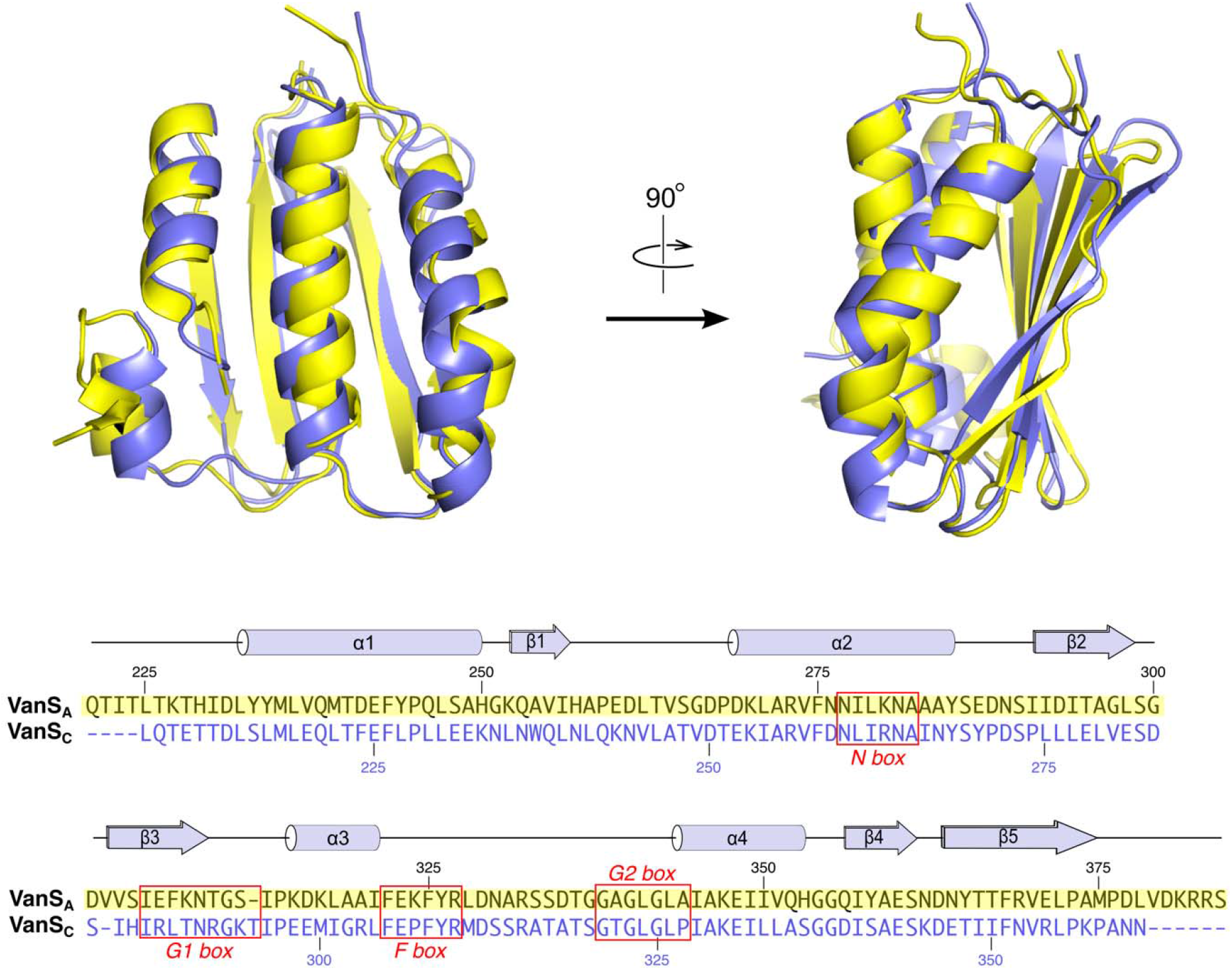
Comparison of the VanS_A_ and VanS_C_ CA domains. Top, superposition of VanS_A_ and VanS_C_, shown in two orthogonal orientations. VanS_A_ is colored yellow, while VanS_C_ is shown in slate blue. Below is shown the pairwise sequence alignment of the two CA domains. Secondary structural elements and conserved sequence motifs are indicated.

Overall, the two structures are very similar. The one exception can be found in the ATP lid; this is not surprising, given the flexible nature that is generally attributed to the ATP lids of CA domains. In particular, helix α3 occupies very different positions in the two structures, being positioned much closer to the nucleotide binding site in VanS_A_ than in VanS_C_ (Figure 2). Notably, α3 is involved in a crystal contact in the VanS_A_ crystals, packing against strand β5 in a neighboring molecule in the lattice (Figure S4). Were α3 to occupy the same position in the VanS_A_ structure that it does in the in VanS_C_ structure, it would clash with the symmetry-related molecule. Hence, it seems likely that the ATP lids of VanS_A_ and VanS_C_ can adopt a range of positions, and crystallization of these two proteins has selected for two different conformers.

### Nucleotide binding

Numerous efforts were made to co-crystallize the VanS_A_ and VanS_C_ CA domains with Mg^2+^ and nucleotide, using both ATP and the non-hydrolyzable analog AMP-PNP. However, no electron density for the nucleotide was ever seen with either protein. This failure to capture the nucleotide-bound forms prompted us to measure the CA domains’ binding affinities for ATP.

We first used the fluorescent ATP analog TNP-ATP. This molecule’s emission profile is highly sensitive to the fluorophore’s environment, and thus it functions as a useful probe for protein binding (25,26). As a positive control, we used the CA domain from *E. coli* EnvZ, for which the TNP-ATP and ATP binding affinities have previously been reported (27). Binding curves were measured for all three CA domains, yielding estimates for TNP-ATP affinity that lie between 40 and 500 μM (Table 2), which all fall within the range reported for TNP-ATP binding by other histidine kinases (27-30). Next, ATP was used to displace TNP-ATP in competition experiments, allowing us to calculate *K*_D_ values for ATP (Figure 3). The ATP binding affinity obtained for the EnvZ CA domain was 290 μM, in reasonable agreement with previously published values, which range from 60-200 μM (27,31). However, the estimated affinities for ATP were much weaker for the two VanS CA domains, falling in the low millimolar range (Table 2). Such *K*_D_ values are atypically high for histidine kinases (32).

**Table 2.** Nucleotide-binding affinities of different VanS constructs.

| **Protein** | **TNP-ATP $K_D$ (μM)** | **ATP $K_D$ (μM)** |
| --- | --- | --- |
| EnvZ CA domain | $43.2 \pm 3.5$ | $290.1 \pm 38.5^a$<br>$83.3 \pm 0.2^b$ |
| VanS<sub>A</sub> CA domain | $499.6 \pm 55.0$ | $10,038 \pm 1785^a$<br>$3948 \pm 1.0^b$ |
| VanS<sub>A</sub> cytosolic domain | $142.6 \pm 6.4$ | $7092 \pm 788^a$ |
| VanS<sub>C</sub> CA domain | $160.2 \pm 10.3$ | $6044 \pm 655^a$<br>$3128 \pm 0.4^b$ |
| VanS<sub>C</sub> N291D CA domain | $870.7 \pm 56.6$ | ND |

<sup>a</sup> Value from TNP-ATP competition experiments
<sup>b</sup> Value from equilibrium dialysis experiments

**Figure 3.**
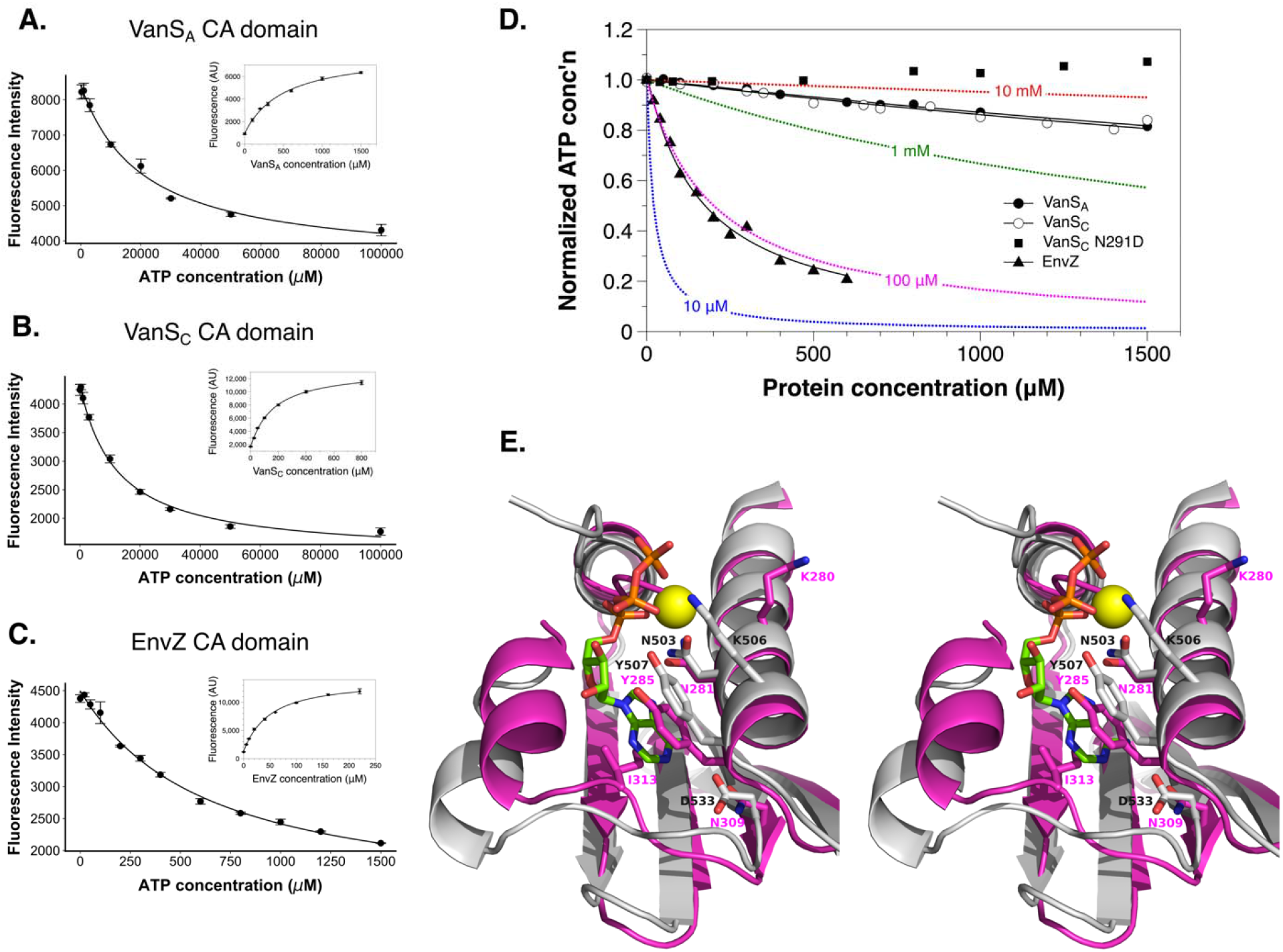
Nucleotide binding by the VanS_A_ and VanS_C_ CA domains. (*A-C*) Binding experiments using TNP-ATP. The insets show direct binding of TNP-ATP, while the main figures show competition assay in which TNP-ATP is displaced by ATP. Lines show fits of the appropriate binding expressions to th data. Panel *A*, VanS_A_ CA domain; panel *B*, VanS_C_ CA domain; panel *C*, EnvZ CA domain. (*D*) Equilibrium dialysis binding experiments with ATP. Solid black lines represent fits of equation 2 to th data. No attempt was made to fit a binding isotherm to the data for the N291D mutant of VanS_C_, since no evidence for binding was seen. Dotted lines in different colors show the theoretical binding curves for selected *K*_D_ values, as calculated from equation 2. Dissociation constants inferred from the binding experiments in this figure are given in Table 2. (*E*) Divergent stereo view of the nucleotide-binding site of VanS_A_ (magenta), superimposed upon the ATP-bound structure of YycG from *B. subtilis* (gray; PDB ID 3SL2). The magnesium ion in the YycG structure is shown as a yellow sphere. Selected amino-acid side chains in and around the nucleotide-binding site are shown.

To ensure that the low affinity for ATP was not an artifact associated with using the isolated CA domain, we repeated this experiment using the entire cytosolic domain of VanS_A_, which includes the HAMP and DHp domains. To prevent ATP consumption, we used the H164A mutant, which lacks the histidine that receives the autophosphorylation. While this cytosolic construct binds TNP-ATP roughly 3-fold more tightly than the isolated CA domain, its affinity for ATP is similar to that of the isolated CA domain (Figure S6A; Table 2). Hence, it does not appear that weak ATP binding can be attributed to use of the isolated CA domains.

We next sought to confirm these results with an orthogonal ATP-binding assay. We chose to use equilibrium dialysis, exploiting the UV absorbance of the nucleotide to monitor free ATP concentration. Again, the *K*_D_ value we obtained for the EnvZ control protein fell within the range of previously published values (27,31), providing confidence in the technique; and again, the binding affinities obtained for the two VanS CA domains were in the low millimolar range, in good agreement with the values obtained from the competition experiments (Figure 3D, Table 2). Therefore, we can say with confidence that the CA domains of VanS_A_ and VanS_C_ proteins have unusually low affinities for ATP, which likely explains our inability to crystallize the protein-nucleotide complexes.

As discussed earlier, the VanS_A_ and VanS_C_ CA domains both contain asparagine in their G1-box sequences, rather than the more commonly occurring aspartic acid. Since this residue forms a salt bridge with the nucleotide’s primary amine in other CA domains, we questioned whether the Asp-to-Asn substitution was responsible for the abnormally low binding affinity of the VanS proteins. To test this idea, we made N309D (VanS_A_) and N291D (VanS_C_) mutants. However, this seemingly conservative change rendered VanS_A_ completely insoluble, both as the isolated CA domain and as the cytosolic-domain construct. The N291D VanS_C_ CA domain was also less stable than its wild-type form, but we were able to express and purify sufficient quantities to perform binding studies. The mutant’s affinity for TNP-ATP was decreased relative to wild-type (Figure S6B); unfortunately, the ATP competition assay could not be performed, as protein precipitated in the presence of high concentrations of ATP. We were able to perform equilibrium dialysis experiments with the mutant, but these assays showed no evidence of ATP binding. Thus, while the interpretation of these experiments is complicated by the instability associated with the Asn-to-Asp mutations, they provide no evidence that restoring an aspartate to the VanS G1-box increases affinity for ATP. Additionally, they suggest that this asparagine plays important roles in the VanS proteins’ folding and/or stability.

Finally, we tested whether VanS prefers GTP over ATP, as has been reported for another histidine kinase with the Asp-to-Asn substitution in its G1 box (33). Using the TNP-ATP competition assay, we compared the relative abilities of GTP and ATP to displace the fluorescent ATP analog, and found that GTP bound less tightly than ATP to both VanS_A_ and VanS_C_ (Figure S7), indicating that it is not the preferred substrate.

### Comparison with CA domains from other histidine kinases

To gain insight into possible factors controlling nucleotide affinity, and to place the VanS_A_ and VanS_C_ CA domains into the broader structural context of prokaryotic histidine kinases, the VanS structures were compared to those of twelve other CA domains. Structures in the comparison set included those of several well-studied CA-domain model proteins, as well as the highest hits identified by BLAST searches using the two VanS CA-domain sequences. Both nucleotide-bound and nucleotide-free structures were included, with three containing no nucleotide, four containing a nucleotide but no associated metal ion, and five containing a nucleotide plus a metal ion (Figure S8). MULTIPROT and SALIGN were used to calculate structural alignments for the fourteen proteins (34,35), with both programs giving essentially identical results. The alignment illustrates that the CA-domain fold is broadly conserved across all proteins considered, with the largest differences being found in the ATP lid (Figure 4).

**Figure 4.**
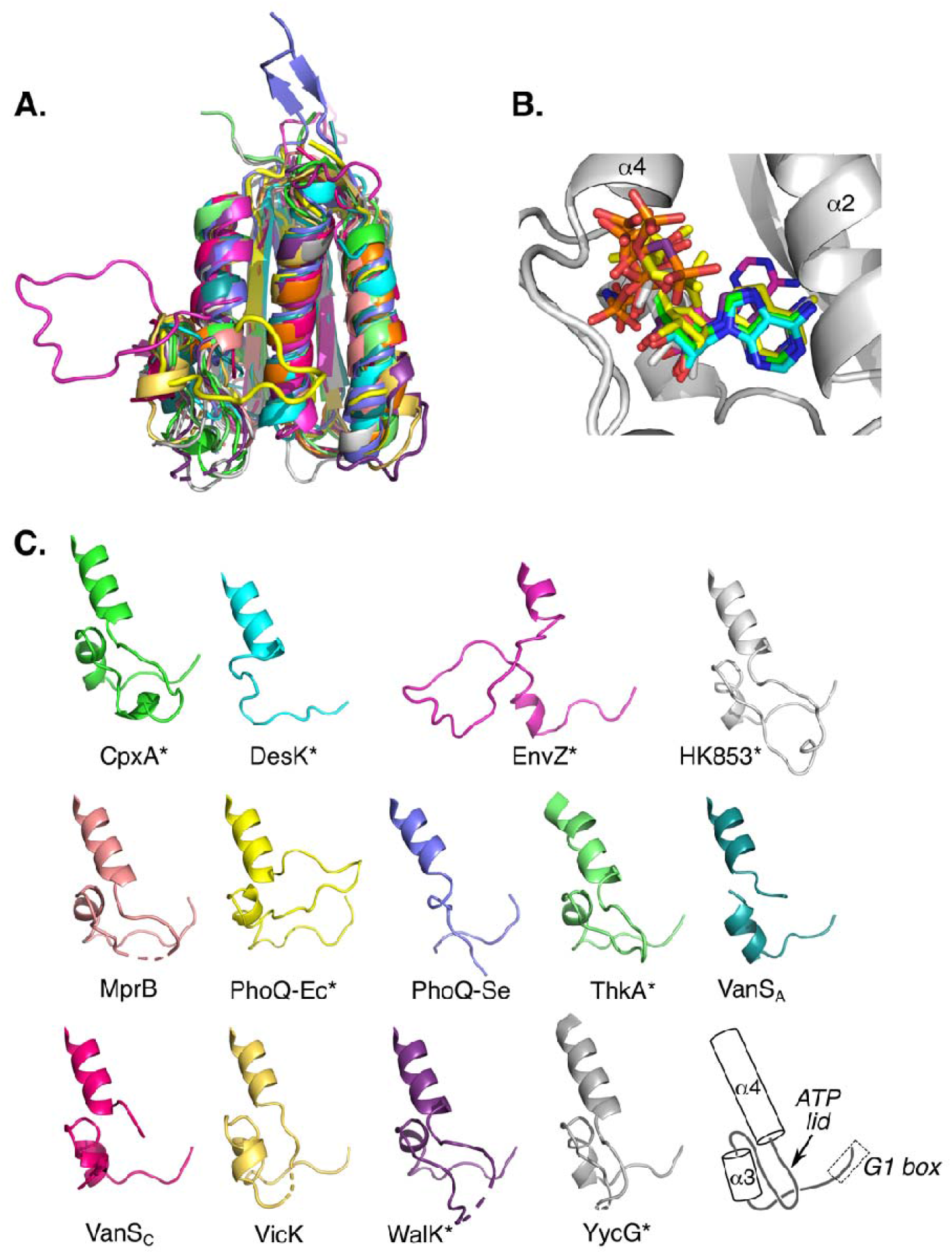
Comparison of VanS_A_ and VanS_C_ with other prokaryotic CA domains. *(A)* Superposition of the VanS_A_ and VanS_C_ CA domains with twelve other CA domains, as performed using the program MULTIPROT (35). Details of the comparison structures can be found in Supporting Information Figure S8. The different proteins adopt highly similar conformations, with the exception of the region between the G1 box and helix 4, which includes the ATP lid. This region lies at the lower left in this panel. (B) Comparison of the nucleotide conformations for the eight structures containing bound nucleotide. A structure cartoon for CpxA (light gray) is included to provide structural context. (C) Comparison of the region between the G1 box and helix α4. All structures were superimposed using MULTIPROT, and are shown in the same orientation; the cartoon at the lower right illustrates the secondary structure elements involved. Color coding in this panel is that same as that used in panel A. Structures containing bound nucleotide are identified with an asterisk. The entire ATP lid is disordered in SrrB, which is therefore omitted from panel (C).

The length of the ATP-lid region differs among the proteins compared, ranging from 39 to 61 residues (for this calculation, the lid is defined as the segment connecting the end of the N-box with the beginning of the G2 box). The corresponding regions in VanS_A_ and VanS_C_ contain 58 and 57 residues, respectively. The largest conformational outlier in the lid region is EnvZ (PDB ID 1BXD), for which the lid forms a loop extending away from the body of the protein. However, EnvZ is the only NMR structure included in the comparison, and its ATP lid was observed to be highly mobile, with a loop conformation that is likely poorly defined owing to a scarcity of medium- and long-range restraints (36). Portions of the ATP lid are also disordered for six of the remaining comparison structures, as well as for VanS_A_ and VanS_C_, all of which contain short regions between helices α3 and α4 that are undefined in the crystal structures. Apart from EnvZ, the other protein with a divergent ATP-lid conformation is *E. coli* PhoQ (1ID0), in which the lid completely covers the nucleotide-binding site, extending from helix α4 all the way to α2. In the remainder of the structures, the ATP lid does not fully bury the nucleotide-binding site, but rather runs alongside of the site, leaving it partially accessible to solvent. In general (and perhaps somewhat surprisingly), there does not appear to be a correlation between the conformation of the ATP lid and the protein’s nucleotide-binding status; for example, *E. coli* PhoQ contains a Mg^2+^-AMPPNP complex and its lid completely buries the bound nucleotide, but other structures that contain Mg^2+^-ATP do not assume the same lid conformation. Furthermore, the presence of a bound nucleotide is not sufficient to lock the ATP lid into a well-defined conformation, as several nucleotide-bound structures still display disorder in portions of their ATP lids (Figure 4).

Helix α3 lies in the middle of the ATP lid region, and its position is roughly similar in all of the compared structures, although in some proteins it is partially unwound (e.g. DesK and *S. typhimurium* PhoQ). VanS_A_ is an exception, as its α3 helix lies closer to the nucleotide-binding site than in any of the other proteins; as described earlier, this likely reflects crystal packing effects.

We next sought to compare the ATP-binding sites of VanS_A_ and VanS_C_ with those of other CA domains, focusing on residues directly involved in binding the nucleotide. First, as noted earlier, most CA domains contain a conserved aspartic acid in the G1 box that forms a salt bridge with an amino group on the adenine base (14); in contrast, the two VanS proteins contain asparagine residues at this position, which adopt the same position and orientation typically associated with the aspartic acid. Therefore, these asparagines should in principle be capable of participating in a hydrogen bond with the base (Figure 3E), and so it is not obvious that this substitution would radically alter nucleotide binding. Secondly, we considered the two hydrophobic residues (Tyr and Ile) that typically sandwich the base, and observed that in the VanS CA domains, these residues are present and occupy positions that are similar to those seen in other CA domains. The same is true for a conserved asparagine in the N-box that helps coordinate the metal ion in the metal-nucleotide complex. Next, we noted that the phosphate groups of the nucleotide typically lie at the N-terminal end of helix α4, where their charge can be partially compensated by the helix dipole. The position of this helix is highly conserved in the various CA domains, including VanS_A_ and VanS_C_, although in the latter two structures, the polypeptide chain N-terminal to the helix actually passes directly through the space that would be occupied by the nucleotide; presumably, this portion of the lid undergoes a rearrangement upon nucleotide binding. Finally, we observed a difference between the VanS proteins and the other CA domains at the C-terminal end of helix α2, immediately downstream of the N-box. In most CA domains, a lysine or arginine is found here, which can form a salt-bridge interaction with the phosphate groups of the nucleotide. In VanS_A_ and VanS_C_, the equivalent position is occupied by an alanine and an asparagine, respectively, neither of which can make a comparable interaction. However, both VanS proteins contain a basic residue one helical turn upstream of this position, within the N-box (Lys-280 and Arg-263 in VanS_A_ and VanS_C_, respectively; see Figure 3E), which may compensate for the loss of the basic residue at the end of helix α2. Overall, then, while the nucleotide-binding sites of VanS_A_ and VanS_C_ differ slightly from those of other CA domains, there are no large structural changes that might explain a significant reduction in ATP affinity.

## Discussion

The VanS proteins play a critical role in the vancomycin-resistance mechanism, by sensing the presence of the antibiotic and, in response, regulating the expression of the resistance phenotype. To expand our understanding of VanS function, we have determined structures for the CA domains of two different VanS proteins, associated with Types A and C VRE. Both proteins adopt the canonical Bergerat fold associated with histidine-kinase CA domains, even though they possess only modest sequence identity with many other CA domains. Our comparison of a panel of different CA domain-containing proteins reveals that all of the proteins studied show strong structural similarity along their entire lengths, with the exception of their ATP-lid regions, which are conformationally diverse.

A surprising result to emerge from this work is the observation that the two VanS CA domains bind ATP with unusually low affinities, with both having *K*_D_ values in the low millimolar range. In contrast, other histidine-kinase CA domains that have been studied bind ATP roughly one to three orders of magnitude more tightly (17,27-31,37). Our affinity estimates are robust, as two orthogonal methods yield *K*_D_ estimates in agreement with each other (Table 2). It is worth noting that our affinity estimate for TNP-ATP binding to EnvZ is 20-fold weaker than previously reported (27); this may reflect the fact that the earlier work titrated TNP-ATP into a fixed concentration of protein, whereas we performed the opposite titration, in order to avoid complications from the inner-filter effect (25). Despite this difference, however, our estimates for EnvZ’s ATP-binding affinity agree well with published values.

The structures of the VanS_A_ and VanS_C_ CA domains do not offer immediate clues to explain their weak binding of ATP. Most of the residues associated with recognition of the nucleotide are either conserved in the two VanS proteins, or can plausibly be replaced by neighboring amino acids. One exception is a highly conserved aspartate normally found in the G1-box, which is replaced by asparagine in both VanS_A_ and VanS_C_. Attempts to produce the Asn-to-Asp mutant yielded unstable proteins in both cases, suggesting that this position may be important for protein folding in ways that are not currently clear. There are regions within the ATP lid that are not well conserved among different CA domains (Figure S8), and it is possible that residues from these regions determine the VanS proteins’ low affinity for ATP.

It is conceivable that low affinities for ATP were observed because the CA domains behave differently in isolation than they do in the context of their full-length proteins. Arguing against this possibility is the observation that the complete cytoplasmic domain of VanS_A_ has a comparable ATP affinity to the isolated CA domain. However, the cytoplasmic-domain construct still lacks the membrane-binding and signal-recognition domains, which could affect nucleotide recognition. On the other hand, CA domains from many other histidine kinases recognize ATP with *K*_D_ values in the low micromolar range, demonstrating that using the isolated CA domains does not necessarily compromise nucleotide binding. Hence, it is plausible that the low affinities we have measured accurately reflect those of the full-length proteins, but this cannot be stated conclusively. Unfortunately, the biophysical assays necessary to directly address this question would require the preparation of many tens of milligrams of purified full-length VanS proteins, which is currently not feasible.

If the low ATP-binding affinities of the CA domains do, in fact, reflect the behavior of the full-length proteins, then the physiological implications are intriguing. Most kinases—including other histidine kinases—bind ATP with affinities in the 1-100 μM range, making the VanS proteins relative outliers (32,38). The VanS proteins’ low affinities mean that their nucleotide-binding sites would essentially never approach saturation in vivo, since bacterial intracellular ATP concentrations typically fall in the 0.5-3 mM range (39-42). This could represent a mechanism that helps minimize basal levels of autophosphorylation occurring in the absence of signal. Alternatively, matching ATP’s *K*_D_ with its cellular concentration could serve to make VanS activity sensitive to changes in ATP concentration, although the utility of regulating vancomycin resistance in this manner is not immediately apparent.

Finally, it is interesting to note that our two CA domains are derived from two different “flavors” of VRE, namely types A and C. Type-A VRE exhibit inducible resistance to vancomycin, i.e. cell wall precursors are not altered in the absence of the antibiotic; in contrast, many (but not all) type-C VRE exhibit constitutive resistance, meaning that the resistance phenotype is observed in both the presence and absence of the antibiotic (9,43). Such a difference would seem to imply a fundamental difference in the regulation of VanS activity between the two VRE types. As both types of VanS protein bind ATP with comparable affinities, it appears that any such difference will not be manifested in the actual autophosphorylation event, but will more likely be found upstream of the ATP-binding or phosphorylation steps.

### Experimental procedures

#### Construct preparation

The VanS_A_, VanS_C_, and EnvZ CA domains were expressed using the pETHSUL vector, which encodes the protein of interest fused with an N-terminally His_6_-tagged SUMO protein (44); the preparation of the cytosolic VanS_A_ construct was described previously (45). The sequence for the VanS_A_ CA domain (residues 221-384) was inserted into the pETHSUL expression vector using ligation-independent cloning, as described (44). The sequences for the VanS_C_ and EnvZ CA domains (residues 208-361 & 289-450, respectively) were inserted into pETHSUL via a DNA assembly reaction using the NEBuilder HiFi DNA assembly kit (New England Biolabs). After cleavage with SUMO hydrolase, the mature VanS_C_ and EnvZ CA domain proteins retain a single vector-derived glycine residue at their N-termini, whereas the VanS_A_ CA domain does not. Primers used for subcloning are shown in Table S1.

#### Expression and purification

The cytosolic VanS_A_ protein was expressed and purified as described (45). Expression constructs for the CA-domain proteins were transformed into BL-21 (DE3) competent cells (New England Biolabs). A single colony was picked and grown overnight at 37°C in LB media containing 100 μg/mL ampicillin (LB-Amp). The overnight culture was diluted 1:50 into 1 L of LB-Amp and placed in a shaking incubator at 30°C. Once the OD reached a value of 0.6-0.8, the temperature was decreased to 20°C and IPTG was added to a final concentration of 0.2 mM. The culture was shaken overnight at 225 rpm and harvested the following morning by centrifugation (3300 x *g*, 40 min). Cell pellets were stored at -80°C before use.

All purification steps were performed at 4°C. Cell pellets were thawed under cold running water and resuspended in Buffer A (50 mM Tris pH 7.8, 300 mM NaCl, 10 mM imidazole, 0.1 mM TCEP) supplemented with 10 mM MgCl_2_ and 2 μg/ml each of DNase and RNase. The cells were lysed by three passes through an Emulsiflex C5 cell disrupter (Avestin, Ottawa, CA). Cell lysates were clarified by sequential low- and high-speed centrifugation steps (15,300 x *g*, 30 mins and 150,000 x *g*, 1 hr, respectively). The clarified lysate was filtered through a 0.45-micron syringe filter and loaded onto a 5-mL IMAC HP column (GE Healthcare) equilibrated with Buffer A. The column was then washed with 20 column volumes of Buffer A and eluted with Buffer B (50 mM Tris pH 7.8, 300 mM NaCl, 250 mM imidazole, 0.1 mM TCEP). To remove the fusion tag, the SUMO hydrolase dtUD1(44) was added to the eluent (1 mg dtUD1/L of culture) and the mixture was dialyzed overnight in Buffer A at 4°C. The following day, the supernatant was passed through a 0.45-micron filter and loaded onto the 5-ml IMAC HP column, re-equilibrated with Buffer A. The flow-through was collected, concentrated with an Amicon Ultra YM-10 concentrator (Millipore), and loaded onto an S100 26/60 size-exclusion column (GE Healthcare) equilibrated in 20 mM Tris pH 7.8, 150 mM NaCl, 0.1mM TCEP and run at 1 mL/min. Both VanS CA domains eluted as single peaks at approximately 80 mins. The purified proteins were concentrated and flash-cooled in liquid nitrogen, and stored at -80°.

#### Mass spectrometry

Proteins were analyzed on a Waters Acquity I-Class UPLC system coupled to a Synapt G2Si HDMS mass spectrometer in positive ion mode with a heated electrospray ionization (ESI) source in a Z-spray configuration. LC separation was performed on a Waters Acquity UPLC Protein BEH C4 1.7 μm 2.1×50mm column. For proteins a 0.2 mL/min gradient of 80/20 to 30/70 A/B in 20 min followed by washing and reconditioning the column was used. Eluent A is 0.1% v/v formic acid in water and B is 0.1% v/v formic acid in ACN. Conditions on the mass spectrometer were as follows: capillary voltage 0.5 kV, sampling cone 40 V, source offset 80 V, source 120 °C, desolvation 250 °C, cone gas 0 L/h, desolvation gas 1000 L/h and nebulizer 6.5 bar. The analyzer was operated in resolution mode and low energy data was collected between 100 and 2000 Da at 0.2 sec scan time. ESI data was deconvoluted using MaxEnt1 in Masslynx 4.1.

#### Crystallization and structure determination

Crystals of the VanS_A_ CA domain were grown using protein at a concentration of 2.5 mg/mL in the size-exclusion buffer described above, supplemented with 2.5 mM AMP-PNP and 5 mM MgCl_2_. Protein and precipitant were mixed in a volume ratio of 1:2 and incubated under Al’s Oil at 18°C (46); the precipitant solution was obtained from the Berkeley Screen (47) and contained 75 mM bis-tris propane, 25 mM citric acid, pH 8.2, 20 mM CdCl_2_, and 25% (w/v) PEG 400. Crystals were taken straight from the drop and flash-cooled in liquid N_2_, with no added cryoprotectant.

Crystals of the VanS_C_ CA domain were grown using protein at a concentration of 2.5 mg/mL in size-exclusion buffer supplemented with 2.5 mM AMP-PNP and 5 mM MgCl_2_. Protein and precipitant were mixed in a volume ratio of 1:3 and incubated under Al’s Oil at 4°C; the precipitant solution was 22% (w/v) PEG 6000, 100 mM buffer. Crystals of comparable appearance and diffraction quality were obtained using MES buffer at pH 6.0, HEPES buffer at pH 7.0, and Tris buffer at pH 8.0; the crystal used for the final data collection was grown at pH 7.0. Crystals were cryoprotected by step-wise transfer from the original mother liquor to a buffer containing 25% (v/v) 2,3-butanediol, 28% (w/v) PEG 6000, 200 mM CaCl_2_, 100 mM HEPES pH 7.0, 2.5 mM AMP-PNP, and 20 mM MgCl_2_. Four steps were used, with approximately three minutes between transfers; after the final transfer, crystals were flash-cooled by plunging into liquid N_2_. Diffraction data were collected at beamlines 24-ID-C of the Advanced Photon Source (APS) and 17-ID-1 of the National Synchrotron Light Source II (NSLS-II); data were processed using XDS (48). The structure of the VanS_A_ CA domain was determined by SAD phasing, using a wavelength of 1.771 Å (7 keV) to exploit the anomalous scattering of the cadmium ions present in the crystallization solution. Using the Autosol pipeline in Phenix, two cadmium positions were found, phases were calculated, and an initial model was built (49). The model was improved by alternating cycles of manual rebuilding in Coot (50) and refinement in Phenix. The initial model was refined separately against the 1.771-Å data set used for phasing and against another data set collected at a wavelength of 0.920 Å (13.5 keV). Both final models contain residues 228-375; no density was observed for residues 323-339 in the ATP lid.

The structure of the VanS_C_ CA domain was determined by molecular replacement. Homology models were built using SWISS-MODEL (51), and the top six models generated in this way were combined using the ENSEMBLER module of Phenix. The resulting probe molecule was positioned using PHASER to generate the initial model, which was subsequently improved by alternating cycles of manual rebuilding in Coot and refinement in Phenix. The final model contains residues 211-359; no density was observed for residues 312-323 in the ATP lid. Data collection and refinement statistics are shown in Table 1. PDB IDs for the structures presented herein are as follows: VanS_A_ CA domain (7 keV data), 8DWZ; VanS_A_ CA domain (13.5 keV data), 8DVQ; VanS_C_ CA domain, 8DX0.

#### Nucleotide binding

In order to measure the binding affinity for TNP-ATP, protein was dialyzed versus 50 mM Tris-Cl, pH 7.5, 50 mM KCl, 5 mM MgCl_2_. Binding reactions were set up in 96-well plates, using a final volume of 100 μL. The TNP-ATP concentration was kept constant at 10 μM, while protein concentration was varied. Fluorescence was measured using a Tecan Spark plate reader fitted with a 510-nm dichroic mirror, using excitation/emission wavelengths of 440 nm and 545 nm, with bandwidths of 15 and 10 nm, respectively. Fluorescence data curves were fit to the following binding expression:

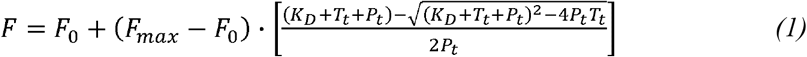

where *T*_*t*_ and *P*_*t*_ are the total concentrations of TNP-ATP and protein, respectively.

For competition experiments, TNP-ATP concentration was kept constant at 10 μM, while the protein concentration was held constant at a value approximately equal to the *K*_d_ for TNP-ATP. Titration series were prepared using a stock solution of 150 mM Mg-ATP in 50 mM Tris-Cl, pH 7.5, 50 mM KCl, 5 mM MgCl_2_ (pH adjusted after nucleotide addition). Fluorescence measurements were conducted as described above. Competition data were fit to the exact competition expression derived by Wang (52).

Equilibrium-dialysis experiments were conducted using a SpectraPor instrument containing five dialysis cells. Each cell consisted of two halves, which were Teflon disks with shallow chambers. The two halves are used to construct a sandwich about a circular piece of 12-14 kDa MWCO dialysis tubing, which separates the two chambers. To prepare for each experiment, protein was dialyzed overnight at 4° versus binding buffer (50 mM Tris pH 7.5, 100 mM KCl, 5 mM MgCl_2_). The five cells were inserted into the instrument, and for each cell one chamber was filled with 300 μL of 140 μM ATP in binding buffer. The other chamber was filled with 300 μL of either binding buffer alone or various concentrations of protein in binding buffer. The five cells were then rotated at 4° for 20-24 hours. Solutions in the ATP side of each cell were then removed and ATP concentrations were determined by absorbance at 260 nM. The buffer absorbance in the buffer-only chamber was also measured, and was invariably found to be equal to that of the opposing, ATP-containing half of the cell, confirming that equilibrium had been achieved. Dissociation constants were obtained by fitting the data to Equation 2 (the derivation of this expression is given in Supporting Information section).

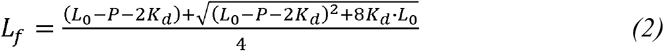

where *L*_0_ = initial concentration of ATP injected into the cell, *L*_f_ = concentration of free ATP at equilibrium, and *P* = the concentration of protein in the cell.

## Acknowledgements

This work was funded in part by a grant from the National Institute of Allergy and Infectious Disease of the National Institutes of Health (R01AI148679). This work includes research conducted at the Northeastern Collaborative Access Team beamlines, which are funded by the National Institute of General Medical Sciences from the National Institutes of Health (P30 GM124165). This research used resources of the Advanced Photon Source, a U.S. Department of Energy (DOE) Office of Science User Facility operated for the DOE Office of Science by Argonne National Laboratory under Contract No. DE-AC02-06CH11357. Support for work performed at the CBMS beam line AMX (17ID-1) at NSLS-II is provided by NIGMS P30GM133893 and DOE-BER KP1607011. NSLS-II is supported by DOE, BES-FWP-PS001. The authors thank Sean Smith for assistance with preparing the VanS_C_ CA domain construct, and gratefully acknowledge helpful discussions with Dr. Yasuyuki Kato-Yamada.

## Conflict of interest statement

The authors declare that they have no conflicts of interest with the contents of this article.

### Abbreviations

VRE: vancomycin-resistant enterococci;
CA domain: catalytic and ATP-binding domain;
DHp domain: dimerization and histidine phosphorylation domain.

## References

1. Levine, D. P. (2006) Vancomycin: a history. Clin Infect Dis 42 Suppl 1, S5–12.

2. Joshi, S., Shallal, A., and Zervos, M. (2021) Vancomycin-Resistant Enterococci: Epidemiology, Infection Prevention, and Control. Infect Dis Clin North Am 35, 953–968.

3. Tacconelli, E., and Magrini, N. (2017) Global Priority List of Antibiotic-Resistant Bacteria to Guide Research, Discovery, and Development of New Antibiotics. World Health Organization

4. Rice, L. B. (2008) Federal funding for the study of antimicrobial resistance in nosocomial pathogens: no ESKAPE. J Infect Dis 197, 1079–1081.

5. Loll, P. J., and Axelsen, P. H. (2000) The structural biology of molecular recognition by vancomycin. Annu Rev Biophys Biomol Struct 29, 265–289.

6. Vollmer, W., Blanot, D., and de Pedro, M. A. (2008) Peptidoglycan structure and architecture. FEMS Microbiol Rev 32, 149–167.

7. Walsh, C. T., Fisher, S. L., Park, I. S., Prahalad, M., and Wu, Z. (1996) Bacterial resistance to vancomycin: five genes and one missing hydrogen bond tell the story. Chem Biol 3, 21–28.

8. Hutchings, M. I., Hong, H. J., and Buttner, M. J. (2006) The vancomycin resistance VanRS two-component signal transduction system of Streptomyces coelicolor. Mol Microbiol 59, 923–935.

9. Guffey, A. A., and Loll, P. J. (2021) Regulation of Resistance in Vancomycin-Resistant Enterococci: The VanRS Two-Component System. Microorganisms 9

10. Capra, E. J., and Laub, M. T. (2012) Evolution of two-component signal transduction systems. Annu Rev Microbiol 66, 325–347.

11. Wuichet, K., Cantwell, B. J., and Zhulin, I. B. (2010) Evolution and phyletic distribution of two-component signal transduction systems. Curr Opin Microbiol 13, 219–225.

12. Zschiedrich, C. P., Keidel, V., and Szurmant, H. (2016) Molecular Mechanisms of Two-Component Signal Transduction. J Mol Biol 428, 3752–3775.

13. Holman, T. R., Wu, Z., Wanner, B. L., and Walsh, C. T. (1994) Identification of the DNA-binding site for the phosphorylated VanR protein required for vancomycin resistance in Enterococcus faecium. Biochemistry 33, 4625–4631.

14. Dutta, R., and Inouye, M. (2000) GHKL, an emergent ATPase/kinase superfamily. Trends Biochem Sci 25, 24–28.

15. Dutta, R., Qin, L., and Inouye, M. (1999) Histidine kinases: diversity of domain organization. Mol Microbiol 34, 633–640.

16. Guarnieri, M. T., Zhang, L., Shen, J., and Zhao, R. (2008) The Hsp90 inhibitor radicicol interacts with the ATP-binding pocket of bacterial sensor kinase PhoQ. J Mol Biol 379, 82–93.

17. Casino, P., Miguel-Romero, L., and Marina, A. (2014) Visualizing autophosphorylation in histidine kinases. Nat Commun 5, 3258.

18. Fihn, C. A., and Carlson, E. E. (2021) Targeting a highly conserved domain in bacterial histidine kinases to generate inhibitors with broad spectrum activity. Curr Opin Microbiol 61, 107–114.

19. Maciunas, L. J., Porter, N., Lee, P. J., Gupta, K., and Loll, P. J. (2021) Structures of full-length VanR from S. coelicolor in both the inactive and activated states. Acta Crystallogr D Biol Crystallogr 77, 1027–1039.

20. Kim, D. J., and Forst, S. (2001) Genomic analysis of the histidine kinase family in bacteria and archaea. Microbiology (Reading) 147, 1197–1212.

21. Yamada, S., Sugimoto, H., Kobayashi, M., Ohno, A., Nakamura, H., and Shiro, Y. (2009) Structure of PAS-linked histidine kinase and the response regulator complex. Structure 17, 1333–1344.

22. Buschiazzo, A., and Trajtenberg, F. (2019) Two-Component Sensing and Regulation: How Do Histidine Kinases Talk with Response Regulators at the Molecular Level? Annu Rev Microbiol 73, 507–528.

23. Zhang, Y., and Skolnick, J. (2005) TM-align: a protein structure alignment algorithm based on the TM-score. Nucleic Acids Res 33, 2302–2309.

24. Kabsch, W. (1976) A solution for the best rotation to relate two sets of vectors. Acta Cryst A32, 922–923.

25. Woodbury, D. J., Whitt, E. C., and Coffman, R. E. (2021) A review of TNP-ATP in protein binding studies: Benefits and pitfalls. Biophysical Reports 1, 100012.

26. Hiratsuka, T. (2003) Fluorescent and colored trinitrophenylated analogs of ATP and GTP. Eur J Biochem 270, 3479–3485.

27. Plesniak, L., Horiuchi, Y., Sem, D., Meinenger, D., Stiles, L., Shaffer, J., Jennings, P. A., and Adams, J. A. (2002) Probing the nucleotide binding domain of the osmoregulator EnvZ using fluorescent nucleotide derivatives. Biochemistry 41, 13876–13882.

28. Guarnieri, M. T., Blagg, B. S., and Zhao, R. (2011) A high-throughput TNP-ATP displacement assay for screening inhibitors of ATP-binding in bacterial histidine kinases. Assay Drug Dev Technol 9, 174–183.

29. Shrivastava, R., Ghosh, A. K., and Das, A. K. (2007) Probing the nucleotide binding and phosphorylation by the histidine kinase of a novel three-protein two-component system from Mycobacterium tuberculosis. FEBS Lett 581, 1903–1909.

30. Sousa, E. H., Gonzalez, G., and Gilles-Gonzalez, M. A. (2005) Oxygen blocks the reaction of the FixL-FixJ complex with ATP but does not influence binding of FixJ or ATP to FixL. Biochemistry 44, 15359–15365.

31. Kenney, L. J. (1997) Kinase activity of EnvZ, an osmoregulatory signal transducing protein of Escherichia coli. Arch Biochem Biophys 346, 303–311.

32. Bhate, M. P., Molnar, K. S., Goulian, M., and DeGrado, W. F. (2015) Signal transduction in histidine kinases: insights from new structures. Structure 23, 981–994.

33. Scaramozzino, F., White, A., Perego, M., and Hoch, J. A. (2009) A unique GTP-dependent sporulation sensor histidine kinase in Bacillus anthracis. J Bacteriol 191, 687–692.

34. Braberg, H., Webb, B. M., Tjioe, E., Pieper, U., Sali, A., and Madhusudhan, M. S. (2012) SALIGN: a web server for alignment of multiple protein sequences and structures. Bioinformatics 28, 2072–2073.

35. Shatsky, M., Nussinov, R., and Wolfson, H. J. (2004) A method for simultaneous alignment of multiple protein structures. Proteins 56, 143–156.

36. Tanaka, T., Saha, S. K., Tomomori, C., Ishima, R., Liu, D., Tong, K. I., Park, H., Dutta, R., Qin, L., Swindells, M. B., Yamazaki, T., Ono, A. M., Kainosho, M., Inouye, M., and Ikura, M. (1998) NMR structure of the histidine kinase domain of the E. coli osmosensor EnvZ. Nature 396, 88–92.

37. Cai, Y., Su, M., Ahmad, A., Hu, X., Sang, J., Kong, L., Chen, X., Wang, C., Shuai, J., and Han, A. (2017) Conformational dynamics of the essential sensor histidine kinase WalK. Acta Crystallogr D Struct Biol 73, 793–803.

38. Becher, I., Savitski, M. M., Savitski, M. F., Hopf, C., Bantscheff, M., and Drewes, G. (2013) Affinity profiling of the cellular kinome for the nucleotide cofactors ATP, ADP, and GTP. ACS Chem Biol 8, 599–607.

39. Bochdansky, A. B., Stouffer, A. N., and Washington, N. N. (2021) Adenosine triphosphate (ATP) as a metric of microbial biomass in aquatic systems: New simplified protocols, laboratory validation, and a reflection on data from the literature. Limnol Oceanogr: Methods 19, 115–131.

40. Buckstein, M. H., He, J., and Rubin, H. (2008) Characterization of nucleotide pools as a function of physiological state in Escherichia coli. J Bacteriol 190, 718–726.

41. Deng, Y., Beahm, D. R., Ionov, S., and Sarpeshkar, R. (2021) Measuring and modeling energy and power consumption in living microbial cells with a synthetic ATP reporter. BMC Biol 19, 101.

42. Yaginuma, H., Kawai, S., Tabata, K. V., Tomiyama, K., Kakizuka, A., Komatsuzaki, T., Noji, H., and Imamura, H. (2014) Diversity in ATP concentrations in a single bacterial cell population revealed by quantitative single-cell imaging. Sci Rep 4, 6522.

43. Sahm, D. F., Free, L., and Handwerger, S. (1995) Inducible and constitutive expression of vanC-1-encoded resistance to vancomycin in Enterococcus gallinarum. Antimicrob Agents Chemother 39, 1480–1484.

44. Weeks, S. D., Drinker, M., and Loll, P. J. (2007) Ligation independent cloning vectors for expression of SUMO fusions. Protein Expr Purif 53, 40–50.

45. Upton, E. C., Maciunas, L. J., and Loll, P. J. (2019) Vancomycin does not affect the enzymatic activities of purified VanSA. PLoS One 14, e0210627.

46. D’Arcy, A., Elmore, C., Stihle, M., and Johnston, J. E. (1996) A novel approach to crystallising proteins under oil. J Cryst Growth 168, 175–180.

47. Pereira, J. H., McAndrew, R. P., Tomaleri, G. P., and Adams, P. D. (2017) Berkeley Screen: a set of 96 solutions for general macromolecular crystallization. J Appl Crystallogr 50, 1352–1358.

48. Kabsch, W. (2010) XDS. Acta Crystallogr D Biol Crystallogr 66, 125–132.

49. Liebschner, D., Afonine, P. V., Baker, M. L., Bunkoczi, G., Chen, V. B., Croll, T. I., Hintze, B., Hung, L. W., Jain, S., McCoy, A. J., Moriarty, N. W., Oeffner, R. D., Poon, B. K., Prisant, M. G., Read, R. J., Richardson, J. S., Richardson, D. C., Sammito, M. D., Sobolev, O. V., Stockwell, D. H., Terwilliger, T. C., Urzhumtsev, A. G., Videau, L. L., Williams, C. J., and Adams, P. D. (2019) Macromolecular structure determination using X-rays, neutrons and electrons: recent developments in Phenix. Acta Crystallogr D Struct Biol 75, 861–877.

50. Emsley, P., Lohkamp, B., Scott, W. G., and Cowtan, K. (2010) Features and development of Coot. Acta Crystallogr D Biol Crystallogr 66, 486–501.

51. Waterhouse, A., Bertoni, M., Bienert, S., Studer, G., Tauriello, G., Gumienny, R., Heer, F. T., de Beer, T. A. P., Rempfer, C., Bordoli, L., Lepore, R., and Schwede, T. (2018) SWISS-MODEL: homology modelling of protein structures and complexes. Nucleic Acids Res 46, W296–W303.

52. Wang, Z. X. (1995) An exact mathematical expression for describing competitive binding of two different ligands to a protein molecule. FEBS Lett 360, 111–114.

53. Diederichs, K., and Karplus, P. A. (1997) Improved R-factors for diffraction data analysis in macromolecular crystallography. Nat Struct Biol 4, 269–275.

54. Weiss, M. S. (2001) Global indicators of X-ray data quality. Journal of Applied Crystallography 34, 130–135.

55. Karplus, P. A., and Diederichs, K. (2012) Linking crystallographic model and data quality. Science 336, 1030–1033.

