## Supporting_information for "Structure of VanS from Vancomycin-Resistant Enterococci: A Sensor Kinase with Weak ATP Binding"

#### *List of Contents*

***Fig. S1.*** Purified proteins used in this work ........ S2
***Fig. S2.*** Representative $VanS_A$ electron density ........ S3
***Fig. S3.*** Representative $VanS_C$ electron density ........ S4
***Fig. S4.*** Crystal packing, $VanS_A$ CA domain ........ S5
***Fig. S5.*** Crystal packing, $VanS_C$ CA domain ........ S6
***Fig. S6.*** Nucleotide binding experiments, cytosolic $VanS_A$ & $VanS_C$ N291D ........ S7
***Fig. S7.*** Comparison of GTP and ATP binding ........ S8
***Fig. S8.*** Sequence alignment of selected CA domains. ........ S9
***Fig. S9.*** Stereo images of the $VanS_A$ and $VanS_C$ CA-domain structures ........ S10
***Table S1.*** Primers used for preparation of expression constructs ........ S11
***Derivation***. Binding expression for equilibrium-dialysis experiments ........ S12
***Supporting Information References*** ........ S13

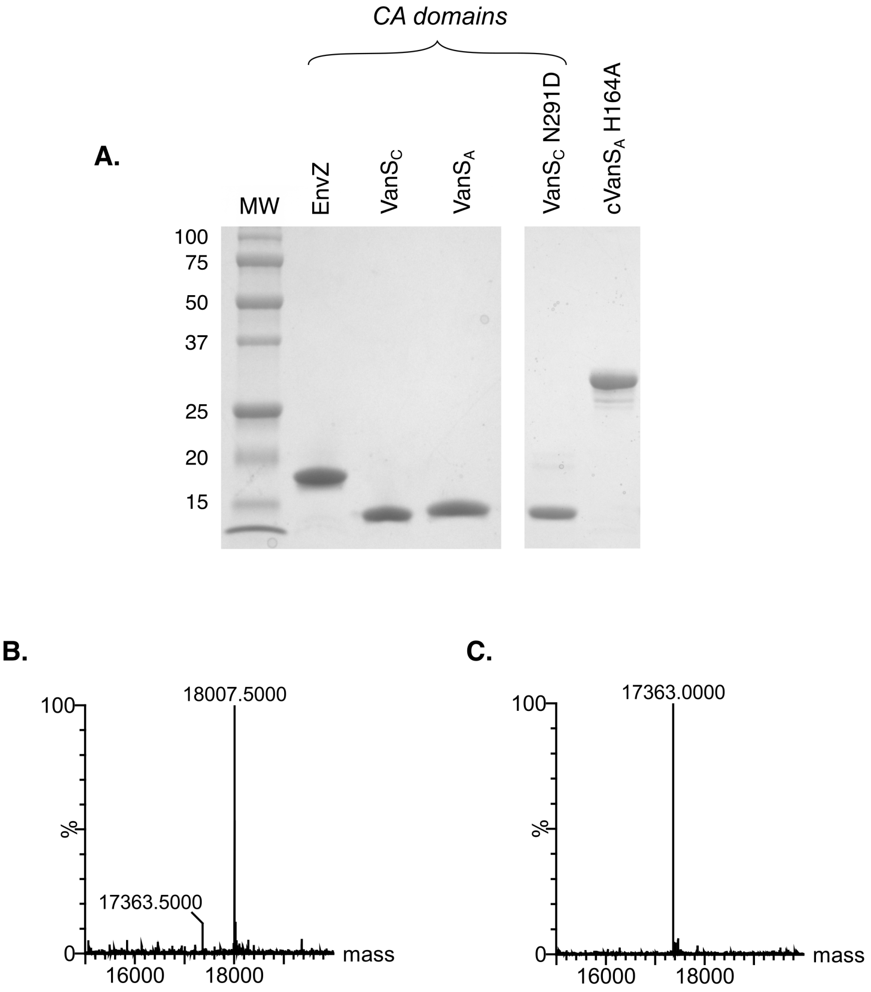


**Figure S1: Purified CA-domain proteins.** (A) Coomassie-stained SDS-PAGE gel showing purified proteins used in this work. The lane labeled "cVanS$_A$" shows the cytosolic domain of VanS$_A$. Note that the VanS CA domains run anomalously on SDS-PAGE gels. (B & C) Deconvoluted mass spectra of the VanS CA-domain proteins, confirming the correct molecular masses. Panel (B) VanS$_A$ (calculated mass 18,008.3); Panel (C) VanS$_C$ (calculated mass 17,363.8). The small peak at 17363.5 Da in panel (B) represents a VanS$_C$ contaminant remaining from an earlier run.

**A.**

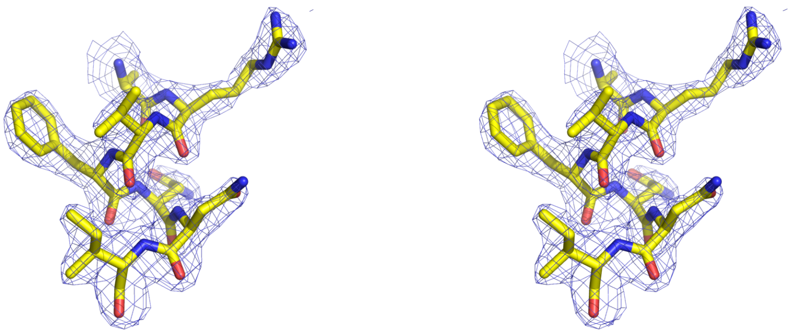

**B.**

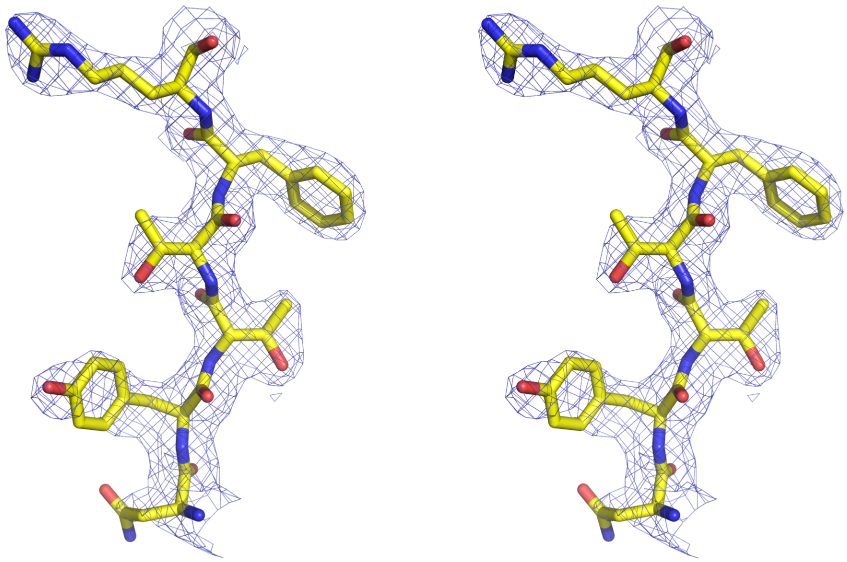

**Figure S2: Representative VanS$_A$ electron density.** Divergent stereo images of 2Fo-Fc electron density, contoured at 1.2σ. (A) Helix α2. (B) Strand β5.

**A.**

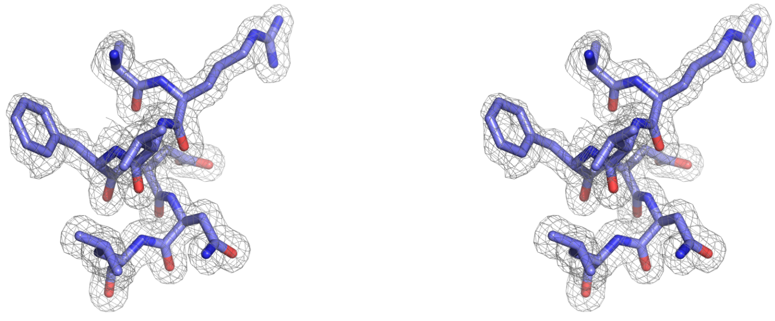

**B.**

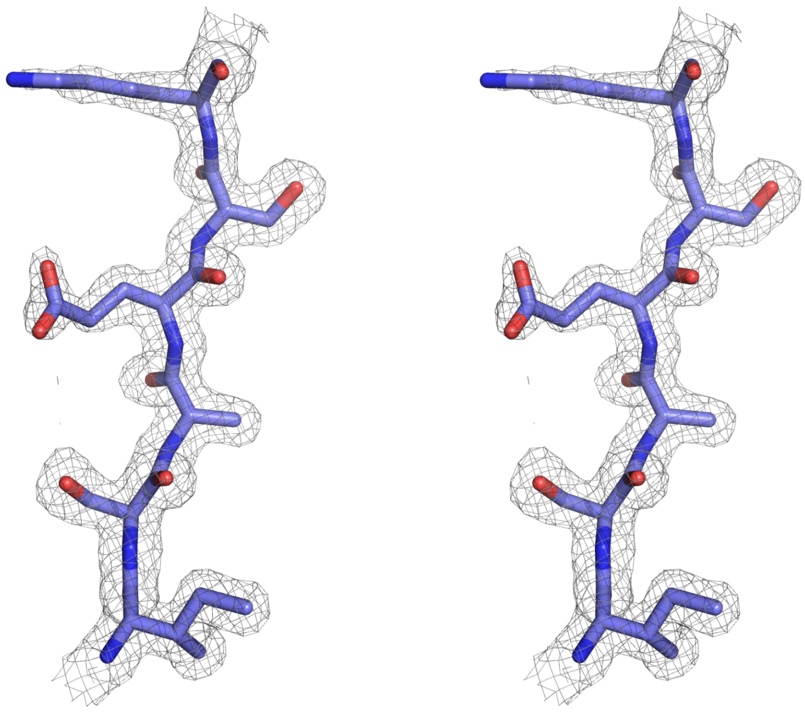

**Figure S3: Representative VanS$_C$ electron density.** Divergent stereo images of 2Fo-Fc electron density, contoured at 1.2σ. (A) Helix α2. (B) Strand β4.

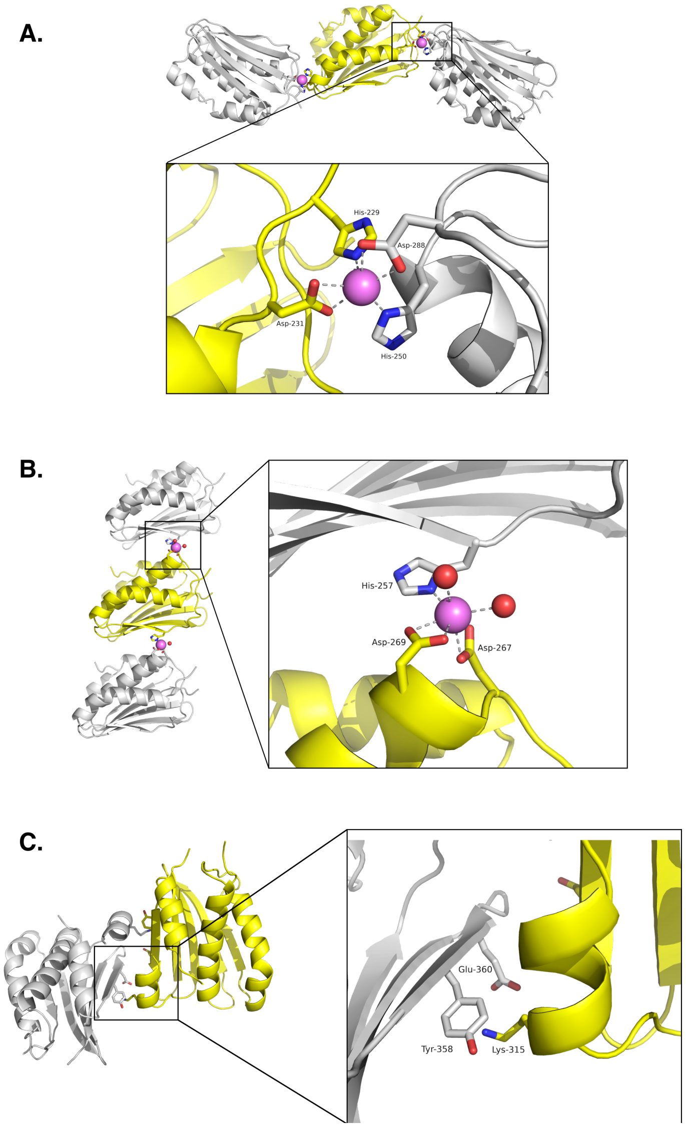


**Figure S4: Lattice contacts in crystals of the VanS$_A$ CA domain.** (A) One of two crystal contacts mediated by solvent-derived cadmium ions. A single VanS$_A$ protein is highlighted in yellow, with two symmetry-related molecules shown in gray; the cadmium ion is shown as a pink sphere. A similar color scheme is used in the other panels of this figure. (B) The second cadmium-mediated crystal contact. Water molecules are shown as small red spheres. (C) Third crystal contact, involving helix $\alpha$3. Lys-315 interacts with either Tyr-358 or Glu-360 on a neighboring molecule (the interaction with Tyr-358 is seen in PDB ID 8DWZ, shown in this panel, while a salt bridge with Glu-360 is seen in 8DVQ). Helix $\alpha$3 sits closer to the main body of the protein in the VanS$_A$ CA domain than it does in other CA-domain structures, likely due to this crystal contact. All three of the lattice contacts shown in this figure are reciprocal, meaning interactions contributed by one molecule are mirrored by corresponding interactions from the symmetry mate; hence, the three types of lattice contacts give rise to six actual contact points/monomer.

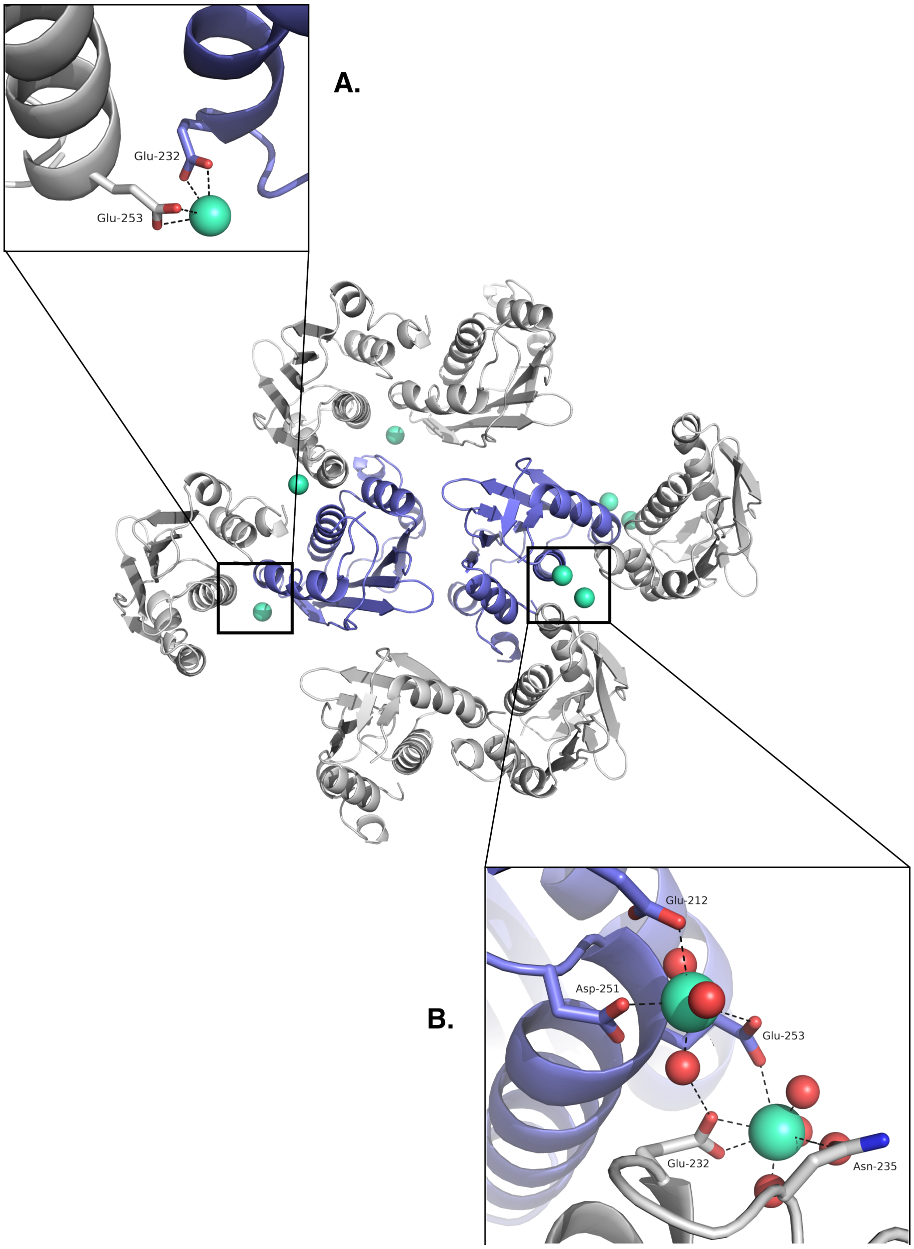


**Figure S5: Lattice contacts in crystals of the $\text{VanS}_\text{C}$ CA domain.** Two distinct magnesium-mediated contacts are seen. The asymmetric unit contains two $\text{VanS}_\text{C}$ chains (shown in blue); selected symmetry-related molecules are shown in gray. Some of the same residues in different chains participate in different contacts. (A) A simple contact involved shared coordination of a single magnesium ion. (B) A more complex contact involving two magnesium ions coordinated between two neighboring chains. Magnesium ions are shown as green spheres, and water molecules are shown as smaller red spheres.

**A.**

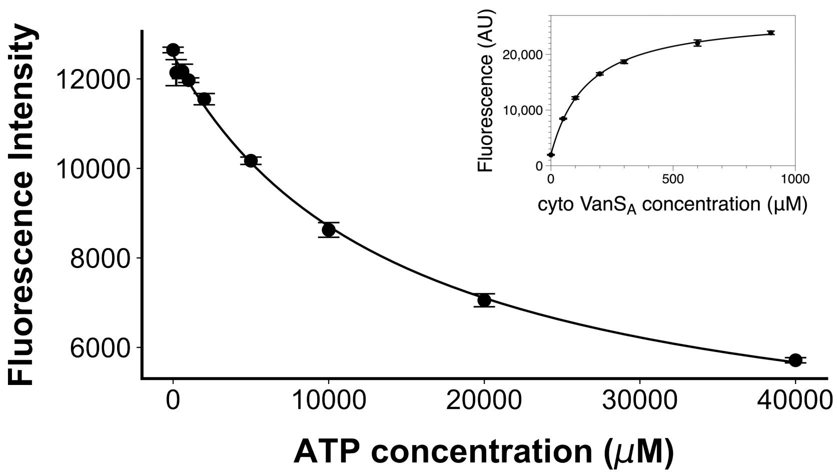


**B.**

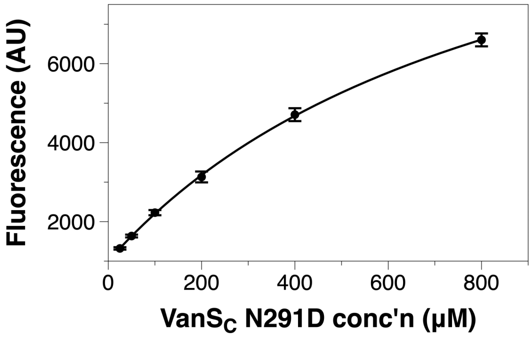


**Figure S6: Nucleotide binding experimental controls.** (A) Binding experiments using TNP-ATP and the cytosolic domain of $VanS_A$. The inset shows direct binding of TNP-ATP, while the main figure shows a competition assay in which TNP-ATP is displaced by ATP. (B) TNP-ATP binding by the N291D mutant of the $VanS_C$ CA domain. Lines show fits of the appropriate binding expressions to the data. Dissociation constants inferred from the binding experiments shown in this figure are given in Table 2.

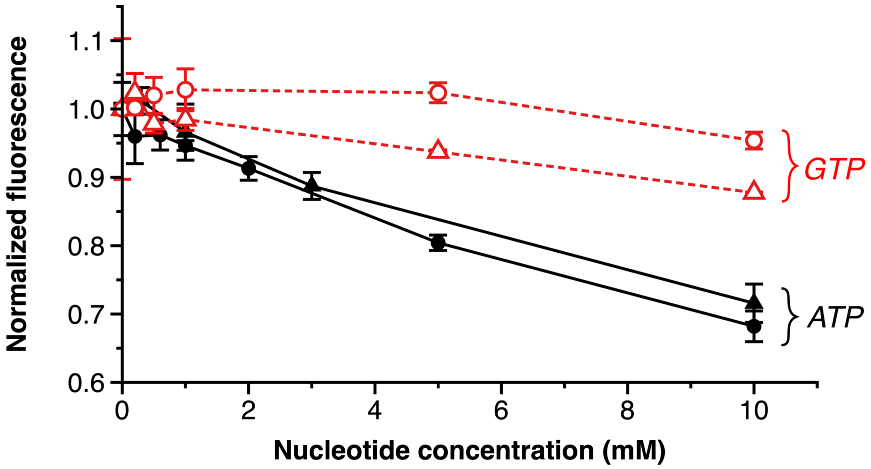


**Figure S7. Comparison of GTP vs. ATP binding.** The TNP-ATP competition assay was used to compare GTP binding with ATP binding. Circles correspond to the cytosolic $VanS_A$ construct, and triangles correspond to the $VanS_C$ CA domain. Filled black symbols represent the competition data obtained with ATP (these data are also shown in Figure 3, panel B, and in Figure S6, panel A); open red symbols show the competition data for GTP. For both constructs, GTP is significantly less effective than ATP at displacing TNP-ATP.

N-box, G1-box

```
VanSA        QTITLTKTHIDLYYMLVQMTDEFYPQ---LSAHGKQAVIHAP---EDLTVSGDPDKLARVFNNILKNAAAYSED---NSIIDITAGL---SGDVVSIEFKNTGS-IPKDK
VanSC        ----LQTETTDLSLMLEQLTFEFLPL---LEEKNLNWQLNLQ---KNVLATVDTEKIARVFDNLIRNAINYSYP---DSPLLLELVE---SDS-IHIRLTNRGKTIPEEM
3A0T ThkA    ----MEFTEFNLNELIREVYVLFEEK---IRKMNIDFCFETD--NEDLRVEADRTRIKQVLINLVQNAIEATGE---NGKIKITSED-MYTK--VRVSVWNSGPPIPEEL
3EHG DesK    -GIRLKDELINIKQILEAADIMFIYE---EEKWPENISL-LN---ENI--------LSMCLKEAVTNVVKHSQA----KTCRVDIQQ--LWKE-VVITVSDDGT------
1BXD EnvZ    TGQEMPMEMADLNAVLGEVIAAESG-----YEREIETALYPG--S--IEVKMHPLSIKRAVANMVVNAARYG-----NGWIKVSSGT-EPNR--AWFQVEDDGPGIAPEQ
3DGE HK385   KSLQINREKVDLCDLVESAVNAIKEF---ASSHNVNVLFESNVPCPVEAY-IDPTRIRQVLLNLLNNGVKYSKKDAPDKYVKVILDE-KDGG--VLIIVEDNGIGIPDHA
1ID0 PhoQ Ec ------RELHPVAPLLDNLTSALNKV---YQRKGVNISLDIS--PE-ISFVGEQNDFVEVMGNVLDNACKYCL-----EFVEISARQ-TDEH--LYIVVEDDGPGIPLSK
3CGZ PhoQ St -SVLLSRELHPVAPLLDNLISALNKV---YQRKGVNISMDIS--PE-ISFVGEQNDFVEVMGNVLDNACKYCL-----EFVEISARQ-TDDH--LHIFVEDDGPGIPHSK
3SL2 YycG    -------MWIQIVRFMSLIIDRFEMT------KEQHVEFIRNLPDRDLYVEIDQDKITQVLDNIISNALKYSPE---GGHVTFSIDV-NEEEELLYISVKDEGIGIPKKD
4I5S VicK    QTSHLDVELTNFTAFMNYILDRFDQI-QSQQSTNKVYEIIRDYPDKSVWIEIDTDKMTQVIDNILNNAIKYSPD---GGKVTITMQT-TDTQ--LILSISDQGLGIPKKD
5C93 WalK    GTTRVDMELVNINEMFNYVLDRFDMILKKDDNPAKYYTIKREFTKRDLWVEIDTDKFTQVLDNIMNNAIKYSPD---GGVVTCRLLE-THNQ--VIISISDQGLGIPRAD
6PAJ SrrB    EGLSVNKEVQPIAALLDKMKIKYRQQ---ADDLGLNMTFNY---CKKRVWSYDMDRMDQVLTNLIDNASRYTK---PGDEIAITCDE-NESE--DILYIKDTGTGIAPEH
4BIV CpxA    KNALVS-ETIKANQLWSEVLDNAAFE---AEQMGKSLTVNFP-PGPWPLY-GNPNALESALENIVRNALRYSH-----TKIEVGFAV-DKDG--ITITVDDDGPGVSPED
6BLK MprB    -GAMVVHEPVDMTEVIDRSLERVRR-----RRSDIEFEVTVT--P--WQVIGDSSGLGRAVLNVLDNAAKWSPP---GGRVGVRLYQIDPGH--AELVITDQGPGIPPQE
```

F-box, G2-box

```
VanSA        LAAIFEKFYRLDNARSSDTGGAGLGLAIAKEIIV-QHGGQIYAESNDNY-TTFRVELPAMPDLVDKRRS---
VanSC        IGRLFEPFYRMDSSRATATSGTGLGLPIAKEILL-ASGGDISAESKDET-IIFNVRLPKPANN---------
3A0T ThkA    KEKIFSPFFTTKT------QGTGLGLSICRKIIEdEHGGKIWTENRENG-VVFIFEIPK--------TPEKR
3EHG DesK    -------FKGEENSFSK-----GHGLLGMRERLE-FANGSLHIDTE-NG-TKLTMAIPNNSK----------
1BXD EnvZ    RKHLFQPFVRGDSART--ISGTGLGLAIVQRIVD-NHNGMLELGTSERGGLSIRAWLPV-PVTRAQGTTKEG
3DGE HK385   KDRIFEQFYRVDSSLTYEVPGTGLGLAITKEIVE-LHGGRIWVESEVGKGSRFFVWIP--KDRAGEDNRQDN
1ID0 PhoQ Ec REVIFDRGQRVDTLR----PGQGVGLAVAREITE-QYEGKIVAGESMLGGARMEVIFGR-----QHSAPKDE
3CGZ PhoQ St RSLVFDRGQRADTLR----PGQGVGLAVAREITE-QYAGQIIASDSLLGGARMEVVFGR-----QHPTQKEE
3SL2 YycG    VEKVFDRFYRVDKARTRKLGGTGLGLAIAKEMVQ-AHGGDIWADSIEGKGTTITFTLPYK--EEQEDDWDEA
4I5S VicK    LPLIFDRFYRVDKARSRAQGGTGLGLAIAKEIVK-QHKGFIWANSEEGEGSTFTIVLPYENDNDAIDEWEED
5C93 WalK    LGHVFDRFFRVDKARSRAQGGTGLGLAISKEVVQ-MLGGRIWVDSVEGKGSTFYISLPYEPYEEE-DLWDDD
6PAJ SrrB    LQQVFDRFYKVDAARTRGKQGTGLGLFICKMIIE-EHGGSIDVKSELGKGTTFIIKLP--KPE---------
4BIV CpxA    REQIFRPFYRTDEARDRESGGTGLGLAIVETAIQ-QHRGWVKAEDSPLGGLRLVIWLP--LYKRS-------
6BLK MprB    RHLVFERFFRSASARS--MPGSGLGLAIVKQVVL-KHGGALRVDYADPAGTAIHIVLPGRPM----------
```

| Protein | Organism | PDB ID | Ligand | RMSD vs. $VanS_A$ (Å) | Sequence identity with $VanS_A$ (%) | RMSD vs. $VanS_C$ (Å) | Sequence identity with $VanS_C$ (%) |
|---|---|---|---|---|---|---|---|
| $VanS_A$ | *Enterococcus faecium* | 8DWZ | none | --- | 100 | 1.68 | 37.4 |
| $VanS_C$ | *Enterococcus gallinarum* | 8DX0 | none | 1.68 | 37.4 | --- | 100 |
| ThkA | *Thermotoga maritima* | 3A0T | Mg-ADP | 2.09 | 24.8 | 1.80 | 22.7 |
| DesK | *Bacillus subtilis* | 3EHG | Mg-ATP | 2.71 | 14.6 | 2.74 | 14.9 |
| EnvZ | *Escherichia coli* | 1BXD | AMPPNP | 2.71 | 21.3 | 2.49 | 22.1 |
| HK385 | *Thermotoga maritima* | 3DGE | ADP | 1.89 | 25.6 | 1.94 | 29.4 |
| PhoQ-Ec | *Escherichia coli* | 1ID0 | Mg-AMPPNP | 2.05 | 22.1 | 2.29 | 20.4 |
| PhoQ-Se | *Salmonella enterica, subsp. enterica, serovar Typhimurium* | 3CGZ | none | 2.30 | 21.4 | 2.40 | 20.1 |
| YycG | *Bacillus subtilis* | 3SL2 | Mg-ATP | 1.94 | 30.5 | 1.83 | 26.6 |
| VicK | *Streptococcus mutans* | 4I5S | none | 2.06 | 33.1 | 1.83 | 29.0 |
| WalK | *Lactiplantibacillus plantarum* | 5C93 | AMPPCP | 2.25 | 29.6 | 1.82 | 26.4 |
| SrrB | *Staphylococcus aureus* | 6PAJ | none | 1.75 | 28.3 | 1.61 | 25.2 |
| CpxA | *Escherichia coli* | 4BIV | ATP | 2.37 | 25.0 | 2.00 | 21.4 |
| MprB | *Mycolicibacterium hassiacum* | 6BLK | Mg-ATP | 1.97 | 22.9 | 1.86 | 20.4 |

**Figure S8.** Top, alignment of the $VanS_A$ and $VanS_C$ CA-domain sequences with those of the twelve homologs shown in Figure 3. Highly conserved residues are shown in red, and moderately conserved residues in blue. The locations of key sequence motifs are indicated. The Asp-to-Asn substitution that occurs in the G1 box of the VanS proteins is highlighted in yellow. Bottom, details of the various homolog structures. Alignment and sequence identities were calculated with COBALT (1); RMSD values were calculated for pairwise alignments using the program TM-align (2).

**A.**

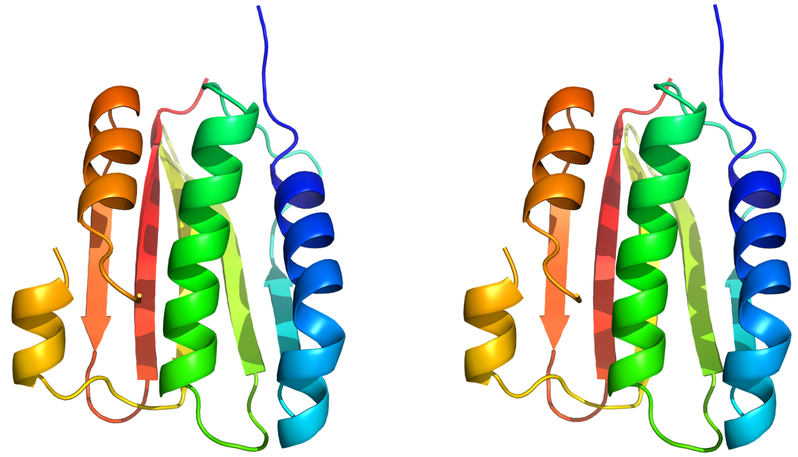

**B.**

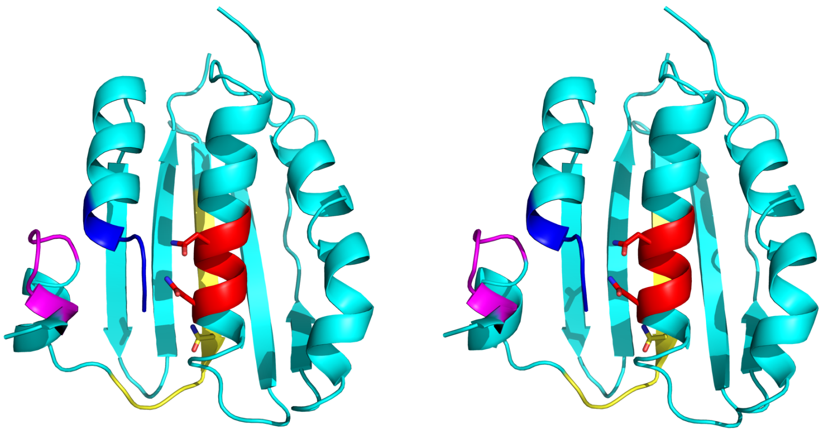

**Figure S9: Stereo images of the VanS CA-domain proteins.** Divergent stereo versions of panels B & C from Figure 1. (A) $VanS_A$ CA domain, shown in a rainbow color scheme (N-terminus = blue; C-terminus = red). (B) $VanS_C$ CA domain, with various functional regions highlighted. N-box, red; G1-box, yellow; F-box, magenta; G2-box, blue.

**Table S1.** Primers used for subcloning of CA-domain constructs.

| Primer | Sequence (5’ to 3’) |
|---|---|
| $VanS_A$ CA domain forward | AGATTGGTGGCCAAACCATTACGCTGACCAAAAC |
| $VanS_A$ CA domain reverse | GAGGAGAGTTTAGACATTAGTCAACCAGATCCGGCATTGC |
| $VanS_C$ CA domain forward | AGATTGGTGGCGGTCTGCAGACGGAAACCACGGAT |
| $VanS_C$ CA domain reverse | GAGGAGAGTTTAGACATTAATTGTTGGCCGGTTTCGGCAG |
| EnvZ CA domain forward | CCGCGAACAGATTGGTGGCGGTCGCACCGGACAGGAGATGCC |
| EnvZ CA domain reverse | CTTCTCGAGGAGAGTTTAGACGATTACCCTTCTTTTGTCGTGCTCTG |

### Derivation of Binding Expression for Equilibrium Dialysis Experiments

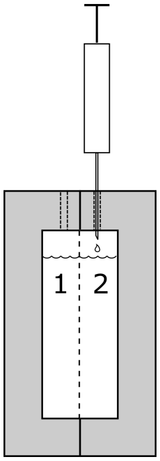


Given a cell with two chambers, separated by a dialysis membrane permeable to ligand but not to protein.

Let $V_1$ = volume of chamber #1, $V_2$ = volume of chamber #2 (for the apparatus used in this paper, the volumes of the two chambers are equal, thus $V_1 = V_2$).

Initiate the experiment by injecting protein (or buffer) into chamber #1 and ligand into chamber #2. After equilibrium is reached, read out free ligand concentration in chamber #2.

$L_0$ = initial concentration of ligand that is injected into chamber #2

$P$ = concentration of protein in chamber #1

$L_f$ = concentration of free ligand at equilibrium

$P_f$ = concentration of free protein at equilibrium

$PL$ = concentration of protein-ligand complex at equilibrium

At equilibrium:

$$L_0 \cdot V_2 = L_f \cdot V_1 + L_f \cdot V_2 + PL \cdot V_1$$ *(Eq. 1: Conservation of mass for ligand)*

Since $V_1 = V_2$, volumes cancel:

$$L_0 = 2L_f + PL$$

$$PL = L_0 - 2L_f$$ *(Eq. 1.1)*

$$P = P_f + PL$$ *(Eq. 2: Conservation of mass for protein)*

*Rearrange Eq. 2 and substitute Eq. 1.1:*

$$P_f = P - PL = P - \left(L_0 - 2L_f\right) = P - L_0 + 2L_f$$ *(Eq. 2.1)*

$$K_d = \frac{P_f \cdot L_f}{PL}$$ *(Eq. 3: Equilibrium expression)*

$$K_d = \frac{\left(P - L_0 + 2L_f\right)\left(L_f\right)}{L_0 - 2L_f}$$ *(Plug Eq. 2.1 into equilibrium expression)*

$$K_d \cdot L_0 - 2K_d \cdot L_f = P \cdot L_f - L_0 \cdot L_f + 2{L_f}^2$$

$$2{L_f}^2 - (L_0 - P - 2K_d)L_f - K_d \cdot L_0 = 0$$

$$L_f = \frac{(L_0 - P - 2K_d) + \sqrt{(L_0 - P - 2K_d)^2 + 8K_d \cdot L_0}}{4}$$ *(positive root is the correct one)*

### References


1. Papadopoulos, J. S., and Agarwala, R. (2007) COBALT: constraint-based alignment tool for multiple protein sequences. *Bioinformatics* **23**, 1073-1079.

2. Zhang, Y., and Skolnick, J. (2005) TM-align: a protein structure alignment algorithm based on the TM-score. *Nucleic Acids Res* **33**, 2302-2309.
